# Direct measurement of B lymphocyte gene expression biomarkers in peripheral blood enables early prediction of seroconversion after vaccination

**DOI:** 10.1101/2020.12.29.424767

**Authors:** Dan Huang, Alex YN Liu, K.S. Leung, Nelson LS Tang

## Abstract

Vaccination is a common and efficient means to reduce the mortality and morbidity of emerging infectious diseases. Among responders, injected antigen induces acquired immunity pathways and leads to the final production of antigen-specific antibodies. The whole process may take weeks to months, depending on the antigen. Typically, seroconversion to influenza vaccine is expected after one month with a responder rate of ~50%.

An early biomarker to predict response is desirable. Peripheral blood gene expression (or transcript abundance, TA) datasets in the public domain were analyzed for early biomarkers among responders. As peripheral blood samples (such as peripheral blood mononuclear cells, PBMC) are cell mixture samples containing various blood cell-types (leukocyte subpopulations, LS). We first develop a model that enables the determination of TA in B lymphocytes of certain genes directly in PBMC samples without the need of prior cell isolation. These genes are called B cell informative genes. Then a ratio of two B cell informative genes (a target gene and a stably expressed reference gene) measured in PBMC was used as a new biomarker to gauge the target gene expression in B lymphocytes. This method having an obvious advantage over conventional methods by eliminating the tedious procedure of cell sorting and enables directly determining TA of a leukocyte subpopulation in cell mixture samples is called *Direct LS-TA method.*

By using a B lymphocyte-specific gene such as TNFRSF17 or TXNDC5 as target genes with either TNFRSF13C or FCRLA as reference genes, the B cell biomarkers were determined directly in PBMC which was highly correlated with TA of target genes in purified B lymphocytes. These Direct LS-TA biomarkers in PBMC increased significantly early after vaccination in both the discovery dataset and a meta-analysis of 7 datasets. Responders had almost a 2-fold higher Direct LS-TA biomarker level of TNFRSF17 (SMD=0.84, 95% CI=0.47-1.21 after log2). And Direct LS-TA biomarkers of TNFRSF17 and TXNDC5 measured at day 7 predict responder with sensitivity values of higher than 0.7. The Area-under curves (AUC) in receiver operation curve (ROC) analysis were over 0.8.

Here, we report a straightforward approach to directly analyses B lymphocyte gene expression in PBMC, which could be used in a routine clinical setting as it avoids the labor-intensive procedures of B lymphocyte isolation. And the method allows the practice of precision medicine in the prediction of vaccination response.

Furthermore, response to vaccination could be predicted as early as on day 7. As vaccination response is based on the similar acquired immunology pathway in the upcoming worldwide vaccination campaign against COVID-19, these biomarkers could also be useful to predict seroconversion for individuals.

## Background

Vaccination by a prior exposure (injection) of an antigen or its precursor is a good strategy for controlling infection and epidemics. Such prior exposure activates the acquired immune system to produce antibody against the pathogen before getting exposed to the pathogen and a full-blown infection. In vaccinated individuals, the pathogens will be controlled quickly, and symptoms of infection are largely reduced or even asymptomatic. A common example is vaccination against the influenza virus. As the prevalent strains of influenza virus change frequently, annual vaccination of different influenza virus strains is commonly practiced in many parts of the World. Of particular relevance is the current pandemic COVID-19 caused by severe acute respiratory syndrome coronavirus 2 (SARS-CoV-2), against which a Worldwide vaccination campaign is also underway.

In the case of vaccination against the influenza virus, just only more than 50% of vaccinated individuals mount an immune response and are protected from subsequent infection. They are called responders (R) to vaccination and are characterized by a having substantial production of antibodies against the antigen injected through the vaccination (seroconversion). On the other hand, the remaining individual with insufficient antibody production and not protected from the virus are called non-responders (NR). Antibody production is measured as titers of antibodies against the vaccination antigen in blood samples taken 28 days after vaccination (Mo et al., 2017). Typically, the reported responder (seroconversion) rate after influenza vaccination is less than 50% of subjects received the vaccination. Furthermore, the responder status will only be known 28 days after vaccination as it takes time for the acquired immune system to produce antibodies against immunization antigen. In order to practice precision medicine of vaccination, and to be better informed of the risk of infection, a better biomarker that can make an early prediction is desirable.

Gene expression profile and changes after vaccination have been studied in various vaccination trials towards different pathogens, including influenza, tuberculosis, hepatitis, and yellow fever (Casey et al., 2019; Henn et al., 2013; Nakaya et al., 2011; Tsang et al., 2014). Most studies measured gene expression or transcript abundance (TA) in a peripheral blood sample, and few of them also did it on purified leukocyte cell-types (Henn et al., 2013). Systemic biology approaches have been applied to study the complexity problem of molecular signatures after vaccination, which is largely confounded by the presence of various leukocyte subpopulations in clinical samples, reviewed by (Pezeshki et al., 2019). Previous studies of TA after vaccination were limited to either focusing on the determinants among responders or the difference between vaccination and a full-blown infection (Rogers et al., 2019). These studies provided a comprehensive list of differential expression genes (DEG) typically composed of tens or hundreds (Henn et al., 2013) genes which were attributed to various pathways (such as interferon or other cytokine response) by gene enrichment analysis. To derive a useful biomarker from a long list of DEG is troublesome both in terms of the applied analysis method has to quantify many genes and lack of clear understanding of the cellular origin of these gene transcripts. Furthermore, most of them used peripheral blood mononuclear cells (PBMC) to study TA in blood samples. Other studies also used whole blood, WB (Weiner et al., 2019). Peripheral blood is composed of various leukocyte cell-types, such as neutrophils, lymphocytes, and monocytes. So it is a kind of convoluted results gathered from many different cell-types present in the blood mixture samples.

Against this background, gene expression (TA) of a single cell-type (subpopulation) is a better biomarker. The research community has called for a new biomarker to enhance vaccine development, particularly in view of the current COVID-19 epidemic(Black et al., 2020; Pezeshki et al., 2019). Here, we are interested if B lymphocyte TA could be a reliable early indicator of subsequent seroconversion as B lymphocyte/plasma cells are the production factory of antibodies, which is the most logical choice of cell-type to follow after immunization. In order to obtain gene expression (TA) level of a single cell-type in peripheral blood, prior isolation and purification of the specified cell-types are required in a conventional approach which could only be carried out in a research laboratory setting as the purification procedure is too tedious to be used in a clinical laboratory. More recently, another method called single-cell RNA-sequencing is also possible to obtain gene expression information of a single cell to obtain the B cell immunoglobulin gene repertoire data (Horns et al., 2020). Single-cell RNA-sequencing (RNA-seq) generates data of gene expression of individual single-cells irrespective of their cell-types by use of expansive equipment and reagents which limits its large-scale application in routine clinical care.

In order to have a better understanding of the cellular origin of a particular gene transcript in commonly used peripheral blood mixture samples, a framework was derived to define a list of cell-type informative genes in a peripheral blood mixture sample, in which the majority (>50%) of the transcripts are contributed by a specific cell-type given both the extent of differential gene expression among those cell-types present in the mixture sample and their respective cell count proportion. The framework of cell-type informative genes is better than the conventional concept of cell-type-specific genes which are only expressed exclusively in a cell-type. Genes with such exclusive expression profiles in the highly related hematopoietic cell-types are very limited (such as *CD19,* CD20 *(MS4A1)* for B cells). Under a similar concept, the latest protein atlas (Uhlen et al., 2019) also has this similar concept and called them as lineage enriched genes which are 50 such B cell lineage genes with the highest expression in the blood (https://www.proteinatlas.org/search/blood_cell_lineage_category_rna%3Ab-cells%3BLineage+enriched+AND+tissue_category_rna%3ABlood%2CLymphoid+tissue%3BIs+highest+expressed+AND+sort_by%3Atissue+specific+score). Here, the two terms, B lymphocytes informative genes and B lymphocyte lineage enriched genes are used interchangeably to indicate the list of genes with higher expression in B lymphocyte in PBMC or whole blood. A similar but not identical method has been used to determine cell count proportions of various cellular subpopulations in a cell mixture by a mathematic model called deconvolution which used a long panel of signature genes for each subpopulation (Avila Cobos et al., 2020; Monaco et al., 2019; Shen-Orr et al., 2010; Shen-Orr & Gaujoux, 2013). However, our target is gene expression (TA) of a subpopulation of interest rather than the proportional count of that subpopulation.

With the list of B lymphocyte informative genes in both PBMC and peripheral blood available, we analyzed their expression profile in publicly available datasets of vaccination experiments. As we are interested in identifying early B lymphocyte TA biomarkers that are predictive of the subsequent antibody response status (seroconversion), we explore the predictive performance of TA data and the Direct LS-TA biomarkers obtained around the first week after vaccination.

## Data, Methods and Analysis Models

A detailed workflow to define B lymphocyte informative genes and Meta-analysis is shown in the supplementary workflow document.

### Datasets used in the analysis of gene expression of peripheral blood and B cells

In order to identify B lymphocyte cell-type informative genes that can predict vaccination response, the following datasets of gene expression obtained from peripheral blood samples were used (Table 1). These datasets were available in the Gene Expression Omnibus (GEO) maintained by the United State National Institute of Health. The details can be obtained under their accession number. The type of blood samples obtained are peripheral blood mononuclear cells (PBMC). In some datasets (GSE45764), further isolation and purification of specific cell-types were performed, such as obtaining the purified B lymphocytes (Henn et al., 2013).

**Table 1.**
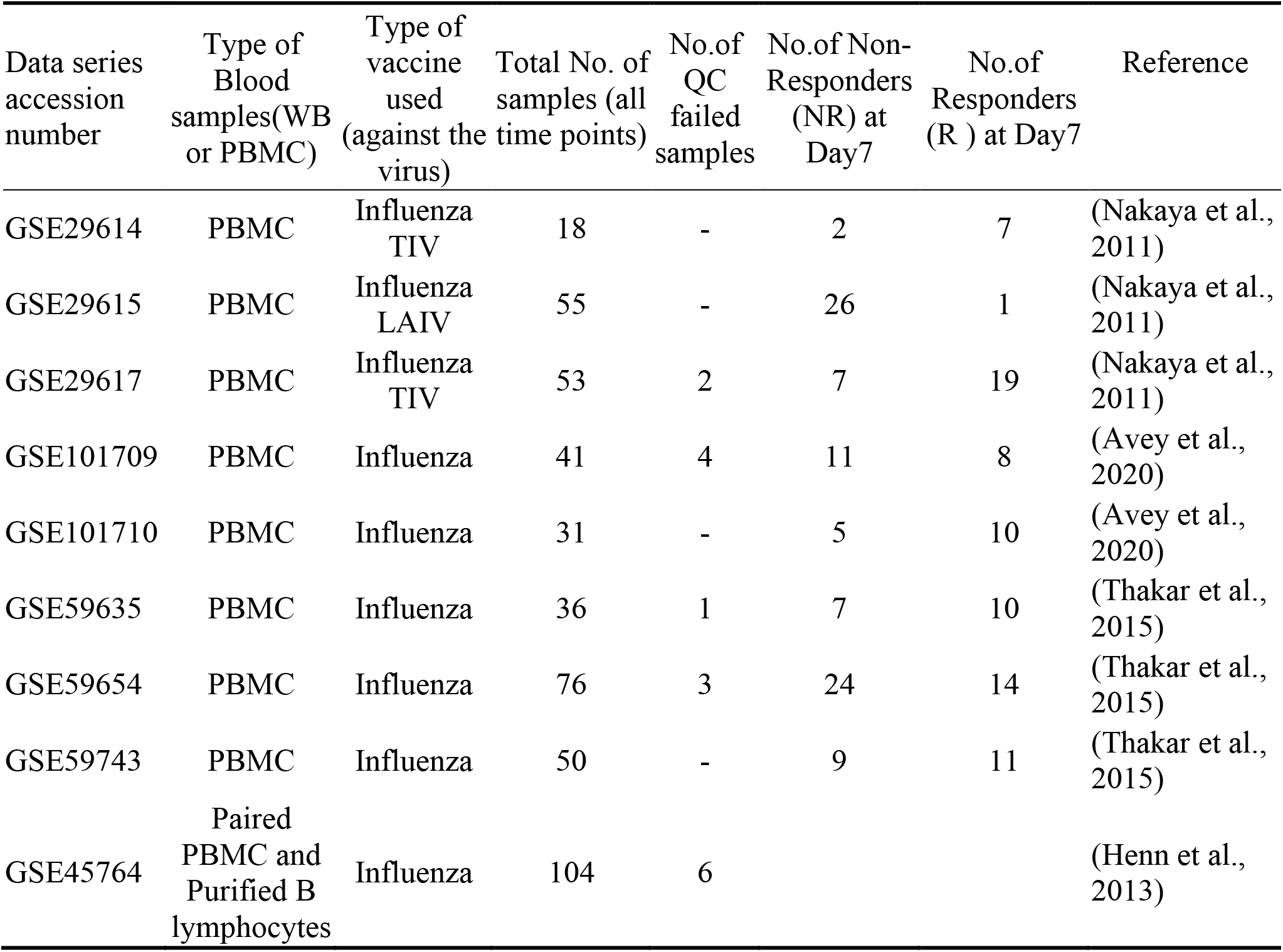
List of gene expression datasets of PBMC or WB used in this study

There are other additional datasets of vaccination studies. However, only these datasets had individual antibody response data, HI titer, available for us to define Responders and Non-responders.

### Definitions of vaccination response

Responders (R) after vaccination are defined following the criteria of seroconversion/significant increase of anti-hemagglutinin antibody levels (HI titer) performed by hemagglutination inhibition (HI) assays on subjects’ plasma or serum has taken before and after vaccination (commonly taken at Day 28)(Mo et al., 2017). The European Committee for Medicinal Products for Human Use (CHMP) defines seroconversion/significant increase as: (a) HI titer after vaccination is at least 1 in 40 and (b) there was at least four folds increase from the pre-vaccination baseline (Committee for Medicinal Products for Human Use., 1997; Mo et al., 2017). Individuals who did not meet these criteria after vaccination were defined as Non-responders (NR).

### To define B lymphocyte informative genes whose expression level can be reliably inferred in cell-mixture samples, e.g., PBMC or WB

A new model was used to incorporate a pre-defined expected proportional count of a particular cell-type (B-lymphocyte in this study) in a cell-mixture sample (e.g., PBMC or WB). Although the B lymphocyte count is variable among individuals and decreases with age (Blanco et al., 2018; Ding et al., 2018), the median proportional cell percentage of B-lymphocyte in WB is about 5%, which is a reasonable pre-defined figure used as an approximation of proportional count in our model (Blanco et al., 2018). Similarly, B lymphocytes are expected to account for ~10% of cells in PBMC.

The model (Figure 1 and 2) is used to shortlist genes that are preferentially expressed by B lymphocytes to the extent such that the sole majority contribution (>50%) of gene transcripts in a cell-mixture sample could be attributed to B lymphocytes. In the example of WB, that is the 5% B lymphocytes component in WB as the sole producer of the majority of gene transcripts (TA) in the WB cell-mixture sample. These genes are called celltype informative genes, in contrast to the conventional concept of cell-type-specific genes which are exclusively produced by a particular cell-type. As transcripts and TA of those informative genes predominantly come from a single cell-type, direct measurement of their total transcript abundance (TA) in the cell mixture sample would be a valid estimation of the gene expression level of that component cell-type.

**Figure 1.**
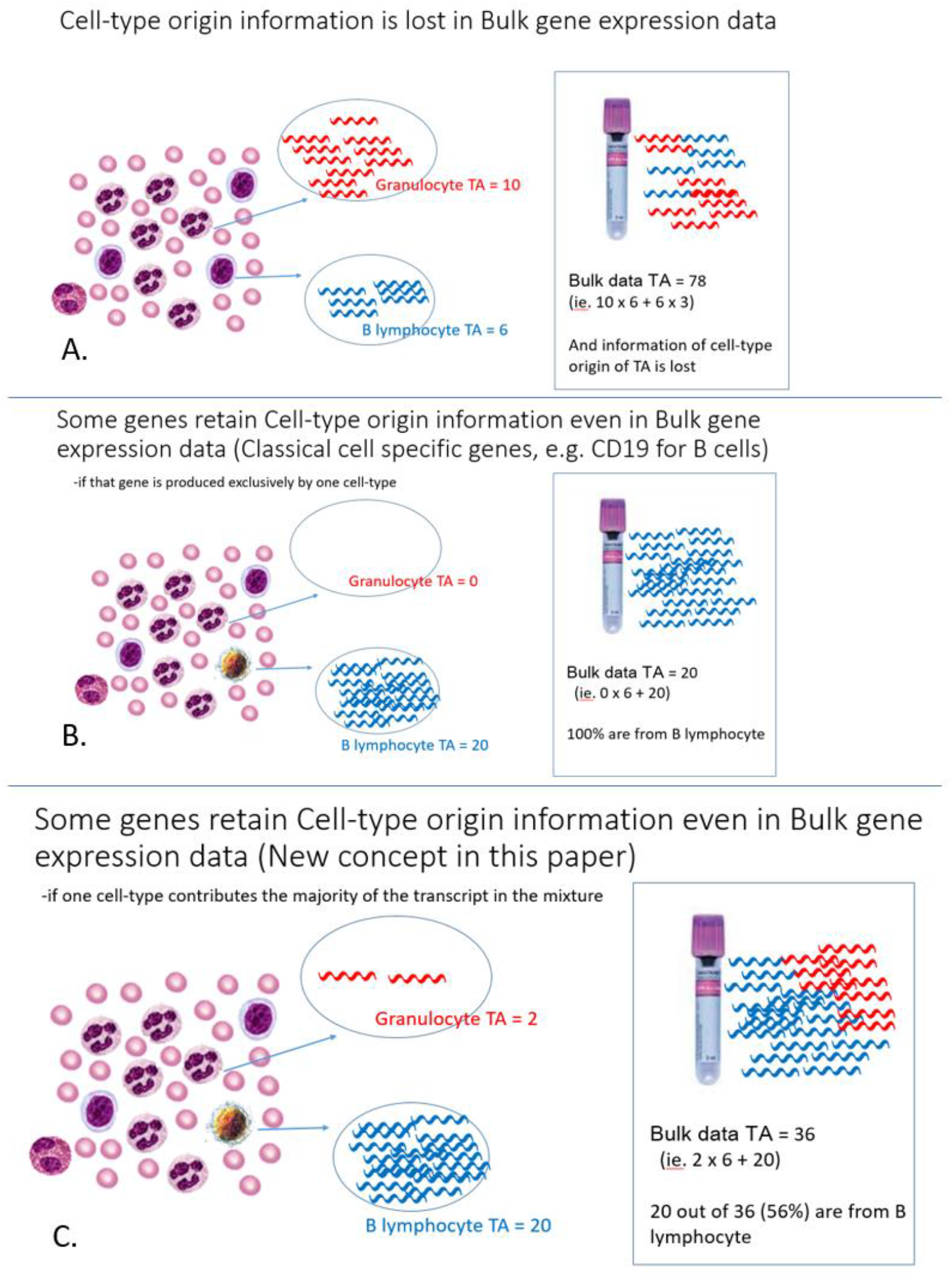
(A) Cell-type origin information is lost in bulk gene expression data. (B) The conventional concept of cell-type-specific genes are those genes exclusively produced by one particular cell-type. Such genes would be few. (C) Our new concept that genes that are preferentially expressed by a cell-type to the extent that it is the sole major contribution (>50%) of gene transcripts in a cell-mixture sample could be informative of gene expression (TA) of that celltype (e.g., B lymphocyte in the figure). Transcript symbols produced by various cell-types are colored for presentation purposes only (Red for granulocyte produced transcripts and blue for B-lymphocyte produced transcripts); the transcripts are, in fact, identical.

**Figure 2.**
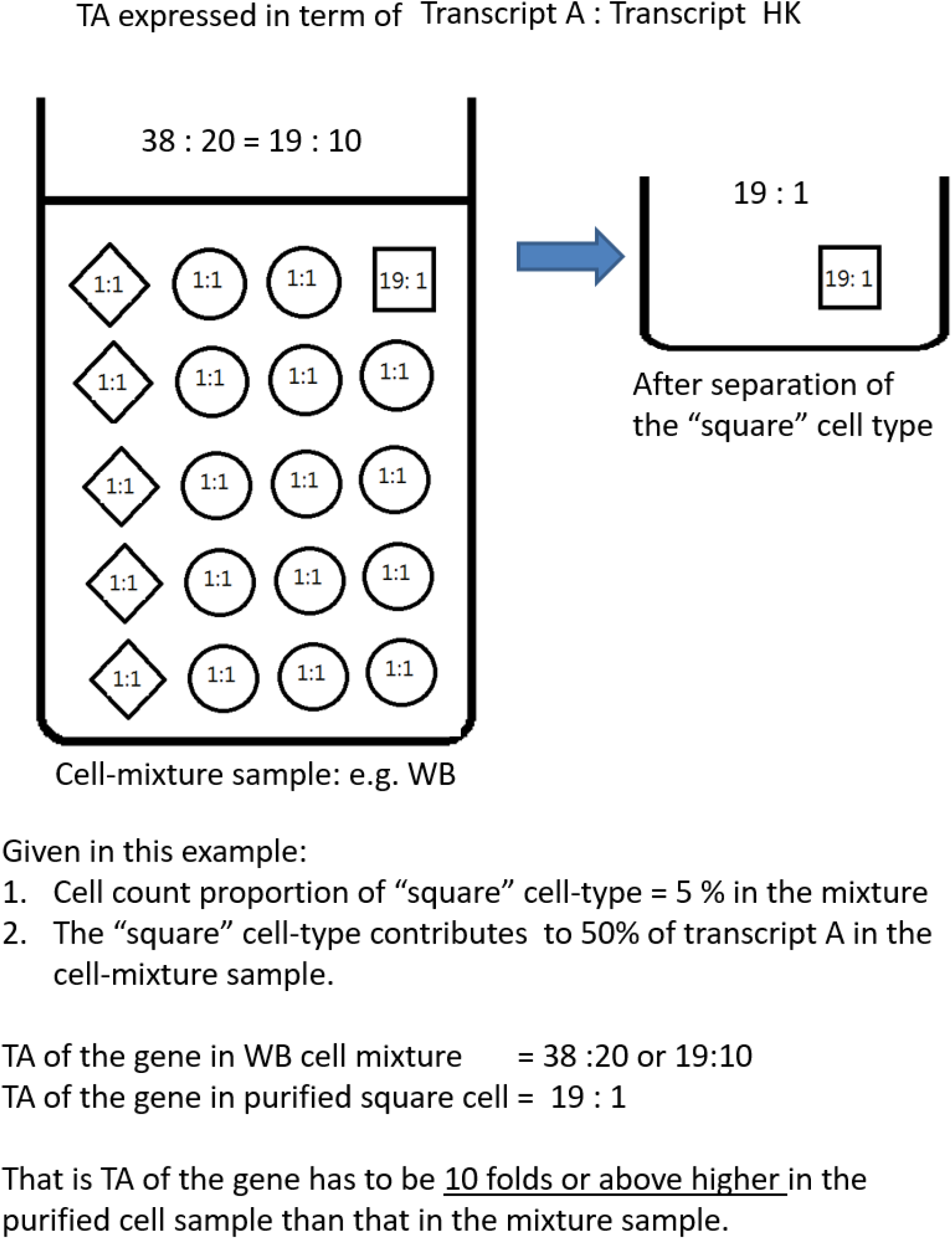
A theoretical model of B lymphocyte cell-type (Square) in a WB sample (with three different cell-types, symbolized as square, circle, and rhomboid). A pre-defined proportional cell count of 5% is given as the proportion of B lymphocyte in WB leukocytes. When TA of a gene was measured both in the WB and purified B lymphocyte samples, B lymphocyte informative genes as we defined will have at least 10-folds higher expression / TA in the purified cell sample (purified B cells) than the cell-mixture sample (WB). Measurement of TA is based on a commonly used normalization approach using another housekeeping (HK) gene.

In order to determine if a gene is a potential B lymphocyte informative gene, it could be evaluated by its TA in both the cell-mixture sample (e.g., WB or PBMC) and purified target cell-type (B lymphocyte) samples. As shown in figure 2, the fold difference between purified B lymphocytes samples and WB samples can be used to determine if a gene in WB had its majority contribution from B lymphocytes. In figure 2, target gene expression is quantified with a housekeeping gene (HK gene), which is a common practice in quantitative PCR. It is shown that any genes having at least 10-folds higher expression in purified B cell samples than WB samples fulfills the criteria of potentially B lymphocyte informative genes. This approach and model is different from previous deconvolution methods of deriving the proportional cell counts of various cell-type subpopulations, the primary output of which are the proportional counts of various cell-types (Monaco et al., 2019; Shen-Orr & Gaujoux, 2013). On the other hand, our method is focused on direct estimation of the average gene expression (TA) of shortlisted informative genes of a given cell-type in WB or PBMC.

### A mathematical model to describe the informative gene framework

In order to understand what would be the required folds difference in TA between purified cells and cellmixture for cell-types of various proportional counts, a mathematical model was used to generalized this concept of the cell-type informative gene to facilitate identification of cell-type informative genes for various cell-types present in different cell-mixture samples (WB, PBMC or other tissue).

Model of % contribution of the transcript by a cell type (e.g., B cells) inside a cell mixture (e.g., WB)

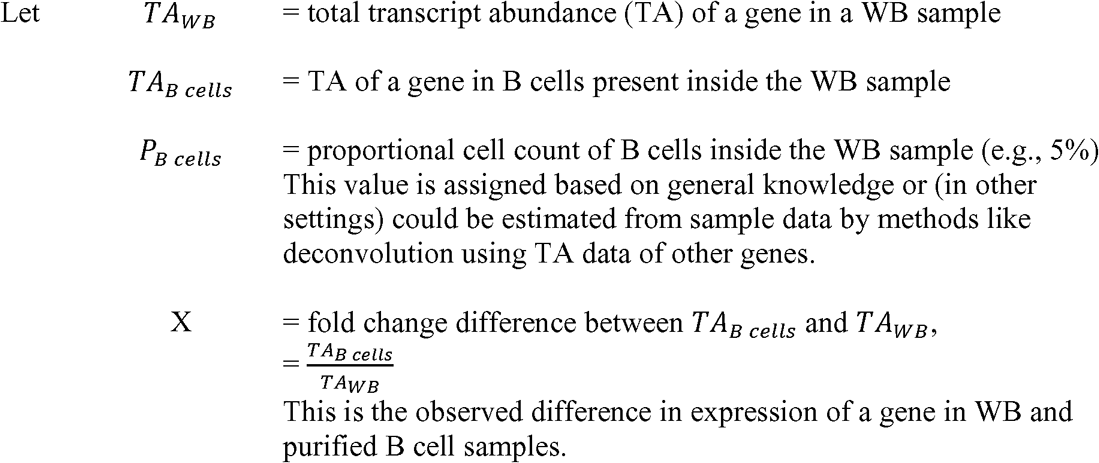

A general model of TA in WB would be

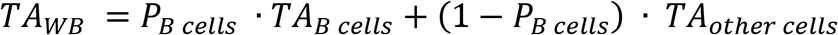

Here *TA_other cells_* is a hypothetical parameter representing a weighted average of TA of all other celltypes based on their respective proportional counts. It is a term that is not actually determined or measured but is used to present a general model.

In order to understand the relationship between % contribution of TA in a cell mixture (WB) by a specified celltype (B cells), which is expressed as Y-axis in supplementary figure 1,

% TA contribution in a WB sample by B cells =

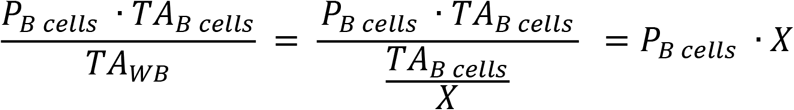

The expression indicates that the relationship between % TA contribution and fold-change difference (X) between specified cell type and WB is linear with the slope defined by the proportional cell count of that cell type.

Supplementary figure 1 shows such a relationship for two scenarios, B cells in WB (given proportional cell count as 5%) and B cells in PBMC (given proportion cell count as 10%). Genes with 10-folds or higher expression (TA) in purified B cells than WB would have B cells contributing at least 50% of transcripts in the WB samples. Similarly, for genes with more than 5-folds or higher expression (TA) in purified B cells than PBMC, at least half of those transcripts in PBMC would originate from B cells. These fold-change criteria are useful to define cell-type informative genes, which could be potentially estimated directly in the cell-mixture sample without the need of prior cell purification.

**Supplementary figure 1.**
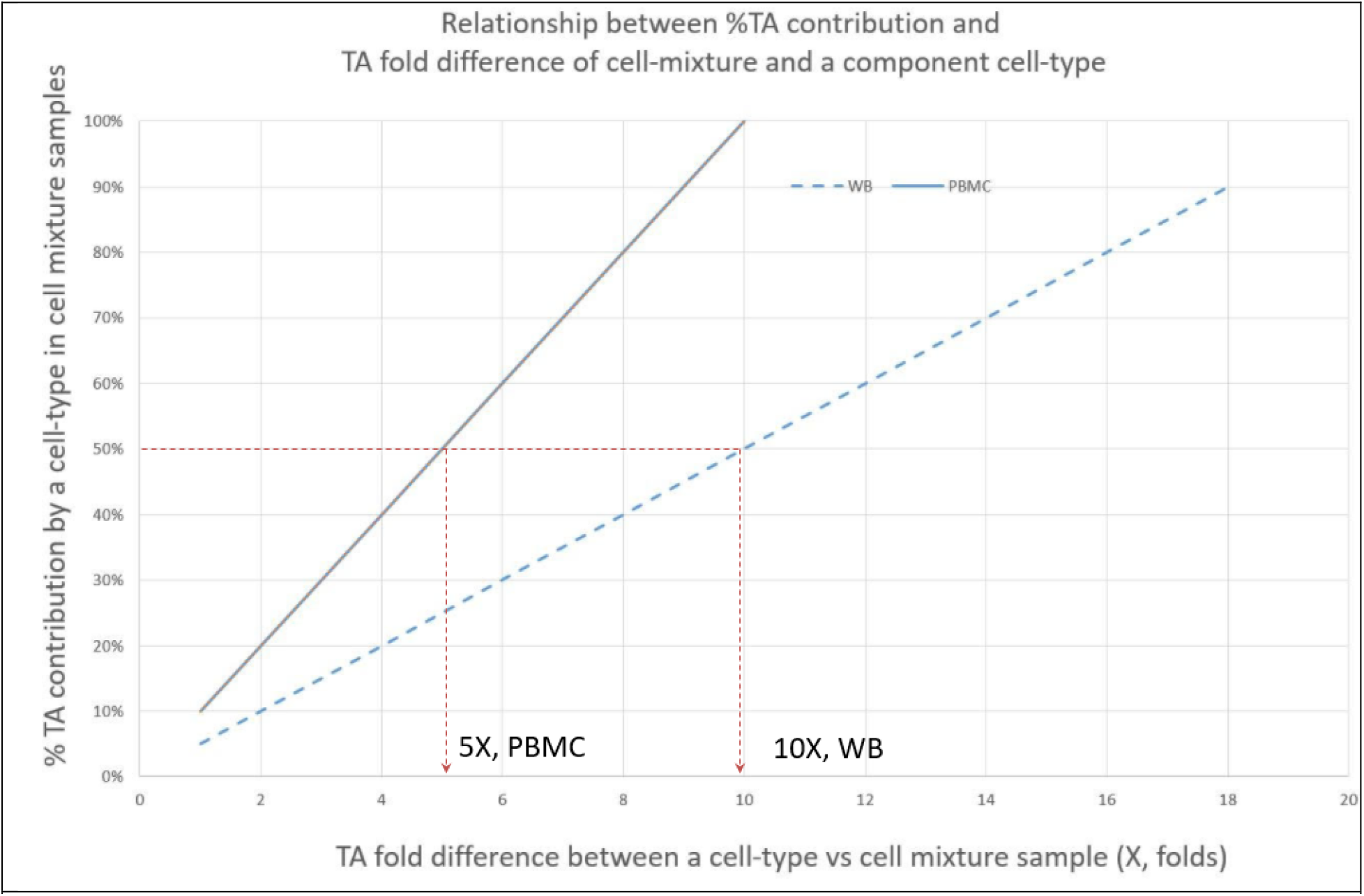
shows such a relationship for two scenarios, B cells in WB (given proportional cell count as 5%) and B cells in PBMC (given proportion cell count as 10%).

An RNAseq dataset (GSE45764) had expression data for samples of purified B lymphocytes with PBMC. Therefore, the datasets were used to identify B lymphocyte informative genes with the given fold-change greater than 5X. Shortlisted potential B lymphocyte informative genes and their biological coefficient of variation (CV%) are shown in supplementary table 1. TA of selected genes were extracted from these datasets for further analysis for their performance as predictors of vaccination response.

#### To identify B lymphocyte reference genes among B lymphocyte informative genes

Biological variation was determined for all B lymphocyte informative genes and expressed in the coefficient of variation % (CV%) in the purified B lymphocyte samples. Those genes with the least CV% were selected as potential B lymphocyte reference genes, which would be used as the denominator in the new biomarker parameter (called Direct LS-TA) to infer B lymphocyte expression directly using the cell-mixture samples (WB or PBMC) data.

#### A new B-lymphocyte biomarker that can be directly measured in whole blood, Direct B lymphocyte LS-TA

Instead of using conventional housekeeping genes (like *GAPDH, UBC)* as the denominator in the expression parameter, another cell-type informative gene has to be used as the denominator in the new Direct LS-TA biomarker for B lymphocytes. It is because conventional housekeeping genes are produced by all the different cell-types in the cell mixture, so it only represents the total cell count of all those various cell-types present in the cell mixture sample but does not represent the variation of cell count of B lymphocyte. Therefore, the normalization factor (or denominator) must also be a cell-type informative gene with an additional feature of having a stable expression which could be measured as CV%.

This new biomarker, Direct B lymphocyte leukocyte subpopulation transcript abundance (LS-TA), is derived from the ratio of TAs of a B lymphocytes target gene and a B lymphocyte reference gene which can be directly quantified from cell-mixture samples without the need to purify B lymphocyte. As log transformation had been applied, this new biomarker parameter was calculated as the difference (subtraction) between the B lymphocyte informative target gene and the B lymphocyte informative reference gene. We focused on two reference genes here (TNFRSF13C and FCRLA), as they had the same low level of CV% with other recognized B lymphocyte-specific genes (e.g., CD19, CD22). Therefore the expression of Direct LS-TA is represented as:

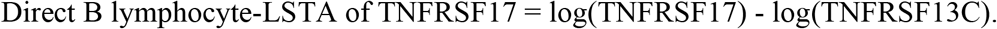

The dataset GSE45764 provided RNA sequencing expression data of paired PBMC and purified B lymphocytes samples taken at the same time. For each potential B lymphocyte informative target genes, they were paired up with the B lymphocyte reference genes (TNFRSF13C and FCRLA) to obtain its Direct B lymphocyte LS-TA results. The ability of Direct B lymphocyte LS-TA results in reflecting the TA of the same target gene in purified B is evaluated by the correlation between Direct B lymphocyte LS-TA measured in the PBMC samples with TA of target genes in the purified B lymphocytes. Pearson’s correlation coefficient (r) was used. A conventional housekeeping gene (e.g., RPL32) was used to control for different numbers of B cells in the purified B cell samples (e.g., X-axis in Figure 3).

**Figure 3.**
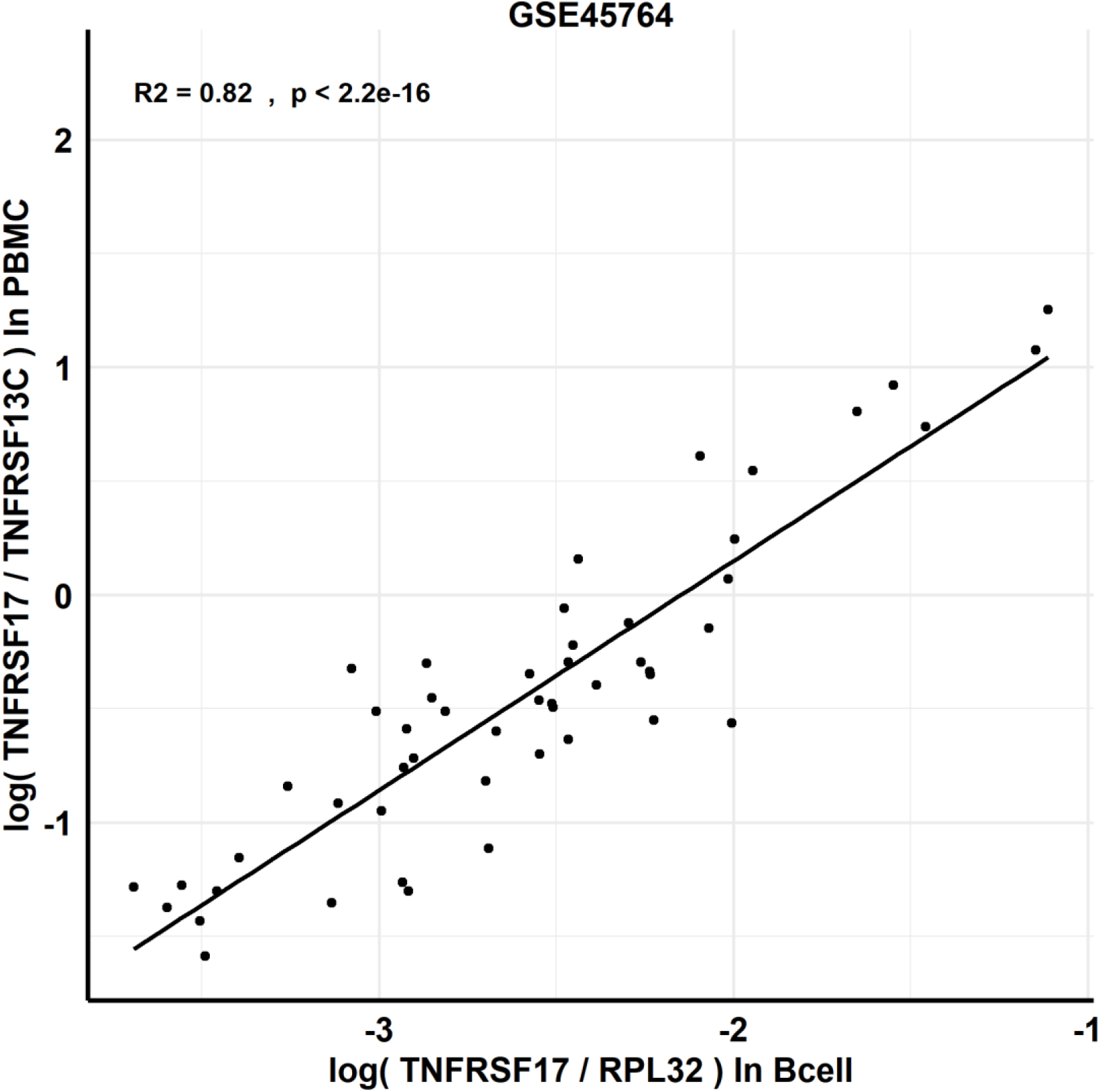
High level of correlation between new biomarker, Direct LS-TA biomarker of target gene TNFRSF17 by measuring TAs of 2 genes (TNFRSF17 and TNFRSF13C) in PBMC samples (Y-axis) and TA of purified B lymphocytes (X-axis). R2=0.82 (p-value < 2.2 × 10^−16)

Using the dataset GSE45764, Pearson’s correlation coefficient (r) of all shortlisted B lymphocyte informative genes were determined, and only those with r > 0.85 were further analyzed for their differential expression between R and NR groups. They represented those B cells target genes that could be reliably inferred from a cell-mixture sample. Thus, they can be efficiently determined in routine clinical samples, and the tedious procedures of cell purification can be eliminated.

In addition, a superseries (GSE59635, GSE59654, GSE59743) contain microarray expression data of paired PMBC and purified lymphocytes samples. They are also used to confirm the list of B lymphocyte informative genes, reference genes, and the performance of Direct B lymphocyte LS-TA of selected target genes.

### Meta-analysis of differentially expressed B lymphocytes LS-TA genes in other datasets

To have a better idea of the reproducibility or replication of these findings, meta-analyses were performed for these 4 Direct B lymphocyte LS-TA biomarkers with PBMC data in a collection of datasets which were performed on different platforms (including Affymetrix and Illumina microarrays).

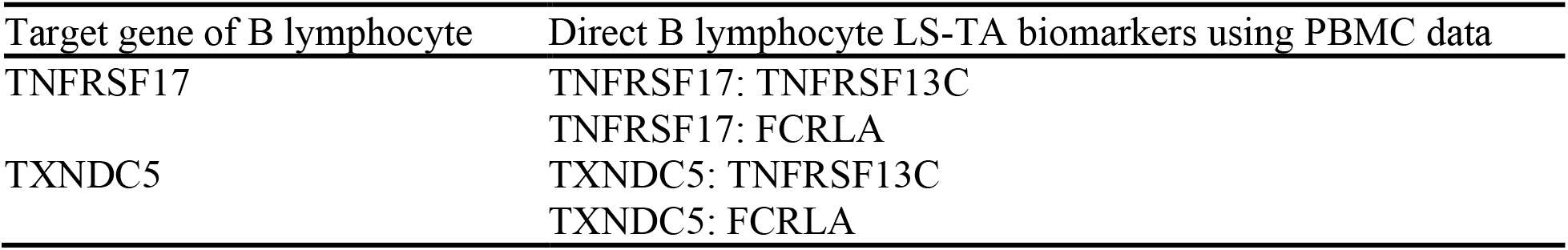

| Target gene of B lymphocyte | Direct B lymphocyte LS-TA biomarkers using PBMC data |
| --- | --- |
| TNFRSF17 | TNFRSF17: TNFRSF13C<br>TNFRSF17: FCRLA |
| TXNDC5 | TXNDC5: TNFRSF13C<br>TXNDC5: FCRLA |

Each dataset was downloaded from GEO. Direct LS-TA results were determined from expression intensities of the target genes and reference genes that had been pickup in the discovery dataset. Fixed effect meta-analysis was performed using R package meta.

### Data analysis and Statistic methods

Microarray datasets were checked if they had been normalized by RMA normalization or quantile normalization. All data were also log-transformed with base 2. Quality check of datasets included a check for outline by Mahalanobis distance metrics (“Mahalanobis Distance,” 2020) using (1) a list of common housekeeping genes and (2) a list of recognized cell-type-specific genes. Samples in a dataset are defined as outliners and removed if they both failed the outliner tests in Mahalanobis distance metrics of (1) and (2). An example of outliner identification in a dataset GSE59654 is shown in supplementary figure 2.

Statistics analyses were performed with R packages, including plotROC, meta, and stats.

**Supplementary figure 2.**
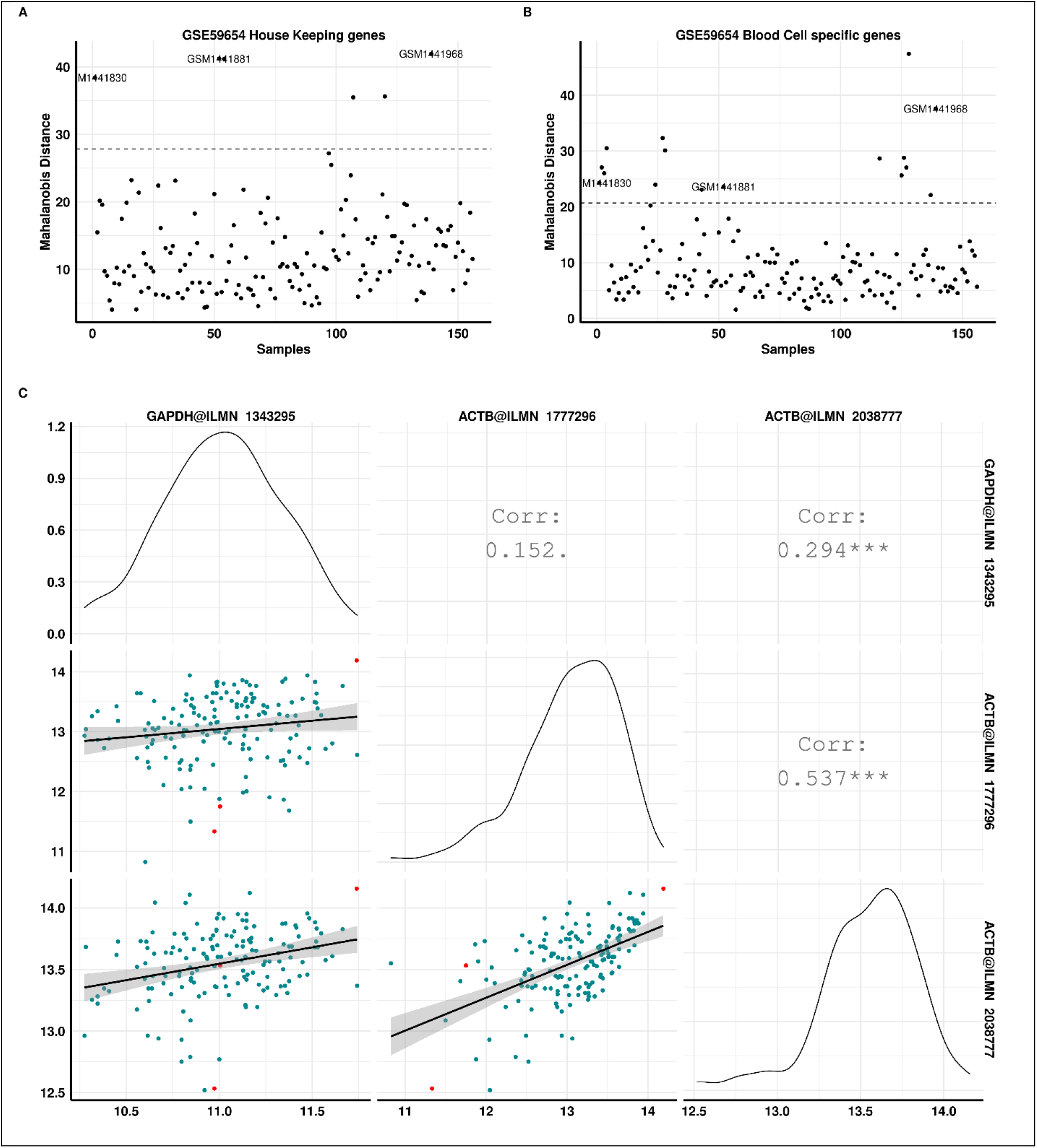
Example of outliner samples in GSE59654, which are labeled as outline by Mahalanobis distance metrics based on (A) a list of housekeeping genes and (B) a list of leukocyte cell-type-specific genes. The three outliner samples called in both (A) and (B) are shown as red symbols on (C) scatter plots of housekeeping genes.

## Results and Discussion

### Shortlisted B lymphocyte informative genes and reference genes

The RNA-seq dataset GSE45764 has both PBMC and purified B cell samples from 5 individuals over multiple days. Given a cell count proportion of 10%, genes expressed more than five-fold higher in purified B lymphocytes than in cell-mixture samples could be used as lymphocyte informative genes. Shortlisted potential B lymphocyte informative genes are shown in Table 2, together with their biological CV% in the purified cell samples. A list of B lymphocyte informative genes with high expression and their biological CV% in PBMC was provided in supplementary table 1.

**Table 2.**
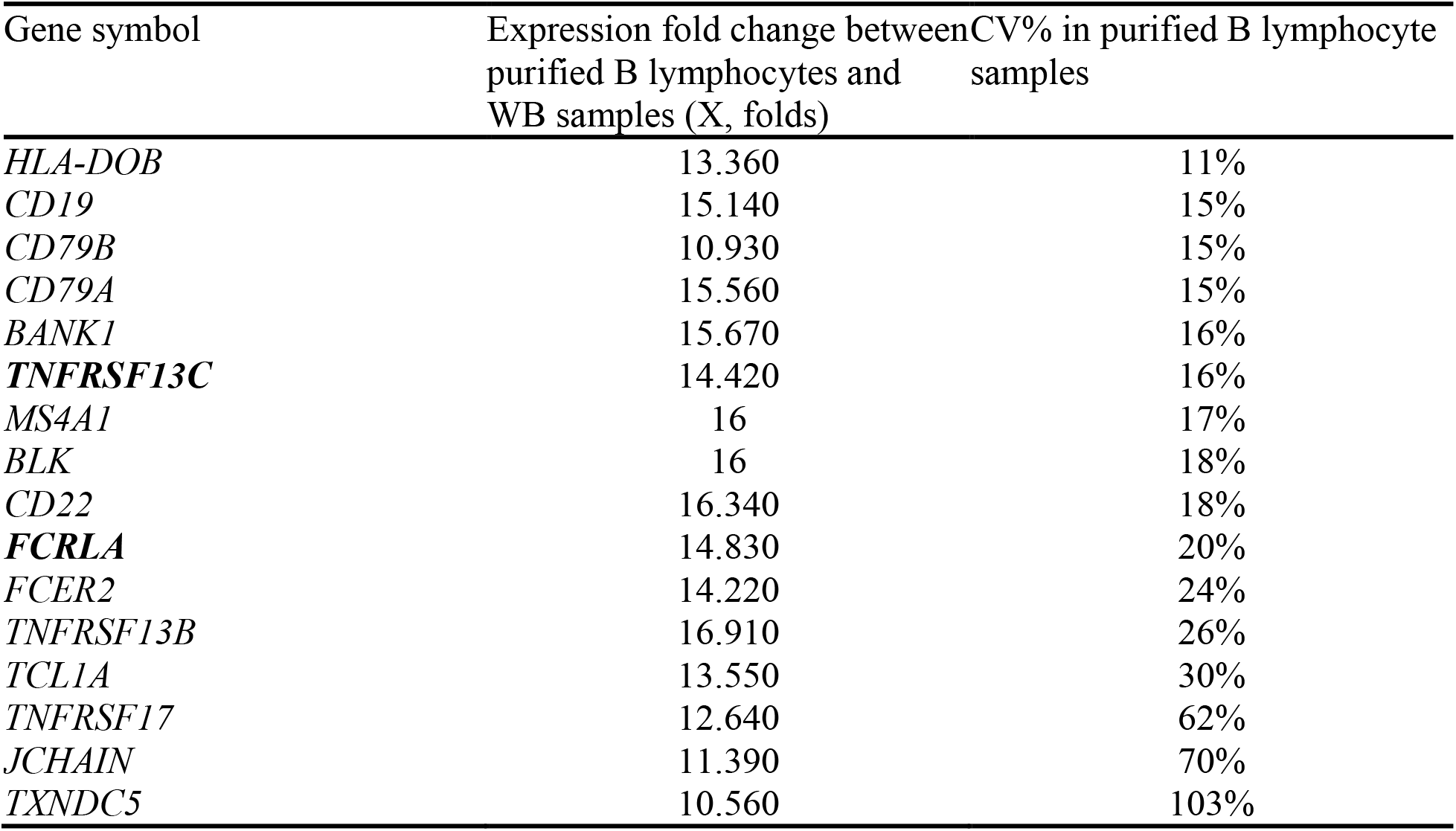
Shortlisted potential B lymphocyte informative genes in RNA-seq dataset GSE45764. Cell-type informative reference genes could be identified among those with low CV%. Two genes stand out among peers of other well-recognized B cell markers, TNFRSF13C and FCRLA, which are further analyzed as B cell informative reference genes in this manuscript.

Among the genes with the least CV% were well-recognized B lymphocyte markers, like CD19, CD20 (MS4A1), and CD22. Interestingly, TNFRSF13C and FCRLA had similar CV% with those recognized B cellspecific markers. Therefore, they were further explored as B lymphocyte reference genes and used as the denominator for the new Direct LS-TA ratio biomarker to predict vaccination response.

### Using Direct determination of B lymphocyte (LS-TA) in PBMC to represent expression (TA) of targeted gene in purified B lymphocytes

In the dataset GSE45764, purified B lymphocytes were collected from 5 individuals over multiple days together with PBMC samples. Therefore, the correlation between new Direct LS-TA biomarkers in PBMC samples could be used to assess its performance to represent expression (TA) of the targeted gene in purified B lymphocytes.

Figure 3 and supplement figure 3 show the performance of the Direct B lymphocyte LS-TA of TNFRSF17 measured in PBMC as a biomarker reflecting TNFRSF17 gene expression in purified B lymphocytes. Direct LS-TA of TNFRSF17 were shown on Y-axis with the expression level of that gene in paired purified B lymphocytes shown on the X-axis. The results confirm that Direct LS-TA TNFRSF17 is a reliable indicator of its expression in purified B lymphocytes. For this example, conventional housekeeping genes were used to normalize results obtained from purified B lymphocyte samples only.

Similarly, Direct LS-TA for another target gene (TXNDC5) in PBMC could represent TA expression levels in purified B lymphocytes. This is shown in both RNA-seq data and microarray data. With such a strong correlation, Direct LS-TA quantification was applied to other vaccination response dataset in which only PBMC samples were collected for gene expression analysis. In fact, the great majority of vaccination response datasets did not have expression information of purified leukocyte subpopulation cell-types; the application of Direct LS-TA method provides a mean to analyze cell-type specific TA in such datasets.

**Supplementary figure 3.**
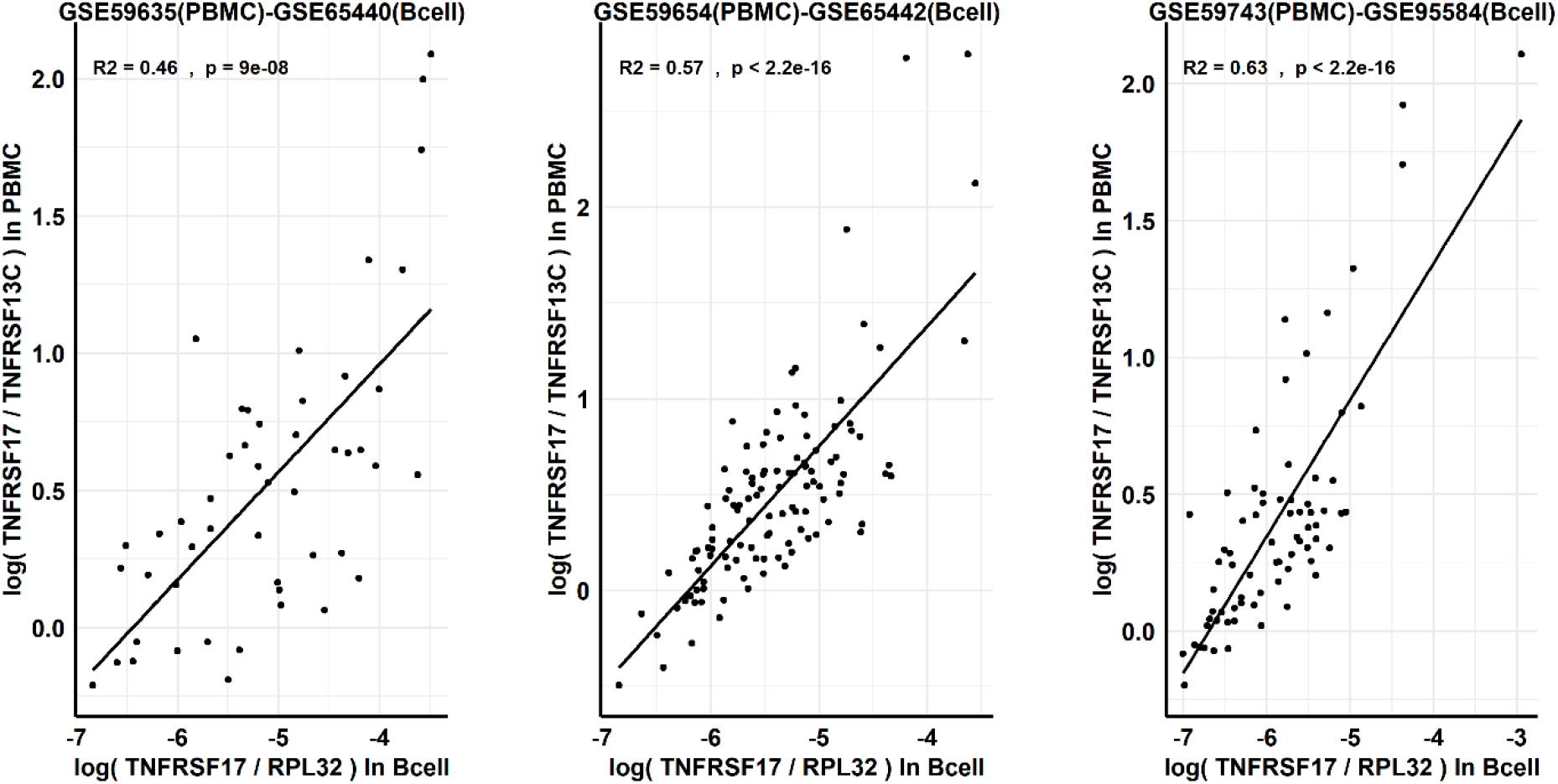
High level of correlation between new biomarker, Direct LS-TA biomarker of target gene TNFRSF17 by measuring TAs of 2 genes (TNFRSF17 and TNFRSF13C) in PBMC samples (Y-axis) and TA of purified B lymphocytes (X-axis).

Not only B-lymphocyte expression of TNFRSF17 could be gauged in a peripheral blood sample, but a list of B-lymphocyte informative genes could also be analyzed using the Direct LS-TA method. Table 3 shows the list of B-lymphocyte target genes whose TA could be estimated in PBMC, and Direct LS-TA biomarker of these target genes had a very good correlation with their TA in purified B lymphocytes. Supplement table 2 also include results using FCRLA as the Direct LS-TA reference gene.

**Table 3.**
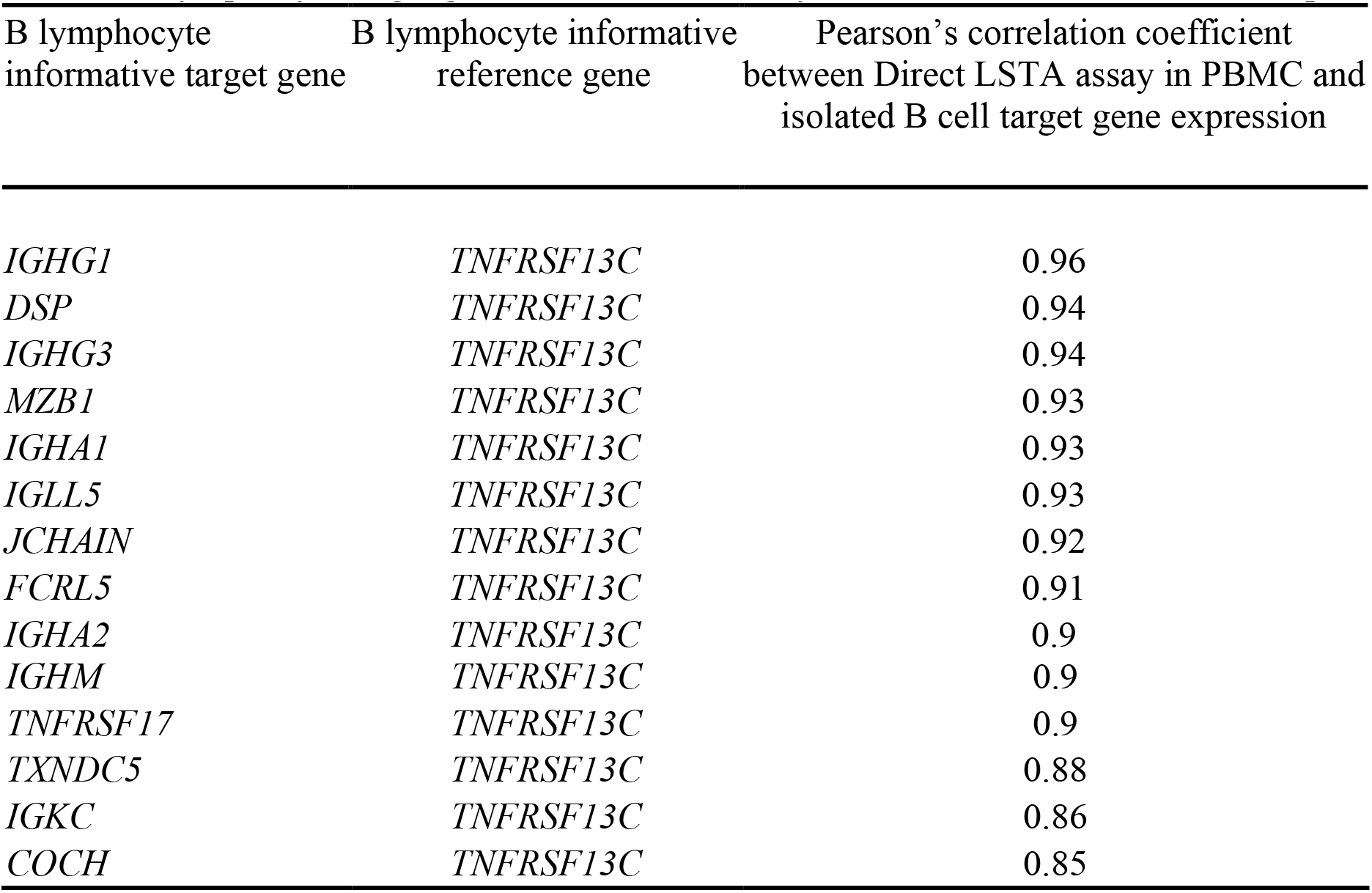
B-lymphocyte target genes that can be reliably measured in PBMC without cell purification

| B lymphocyte informative target gene | B lymphocyte informative reference gene | Pearson's correlation coefficient between Direct LSTA assay in PBMC and isolated B cell target gene expression |
| --- | --- | --- |
| *IGHG1* | *TNFRSF13C* | 0.96 |
| *DSP* | *TNFRSF13C* | 0.94 |
| *IGHG3* | *TNFRSF13C* | 0.94 |
| *MZB1* | *TNFRSF13C* | 0.93 |
| *IGHA1* | *TNFRSF13C* | 0.93 |
| *IGLL5* | *TNFRSF13C* | 0.93 |
| *JCHAIN* | *TNFRSF13C* | 0.92 |
| *FCRL5* | *TNFRSF13C* | 0.91 |
| *IGHA2* | *TNFRSF13C* | 0.9 |
| *IGHM* | *TNFRSF13C* | 0.9 |
| *TNFRSF17* | *TNFRSF13C* | 0.9 |
| *TXNDC5* | *TNFRSF13C* | 0.88 |
| *IGKC* | *TNFRSF13C* | 0.86 |
| *COCH* | *TNFRSF13C* | 0.85 |

Many target genes encode for part of the antibody (like *IGHG1, IGLL5).* Other genes were *MZB1, FCRL5, TNFRSF17, and TXNDC5* which were described in B lymphocyte. The function of 2 other genes, *DSP and COCH,* was not certain.

### Differential expression of B lymphocyte LS-TA at first week after vaccination

With this new B lymphocyte biomarker available, it is now possible to estimate the expression of B lymphocyte as a response to vaccination using the expression data of PBMC samples (see the list in table 1). Dataset GSE59654 (influenza vaccine) was used as an explorative dataset to visualize any differential gene expression (as determined by LS-TA) at Day 7 between the two groups of R and NR. This dataset was selected as it had the largest sample size. Figure 4A shows that R had a significantly higher level of Direct B lymphocyte LS-TA of TNFRSF17 than NR (Wilcoxon test, p-value =0.037). Similarly, the increment of Direct B lymphocyte LS-TA of TNFRSF17 from the day of vaccination (Day 0) to Day 7 was also highly significant in a paired Wilcoxon test (Figure 4B, p-value =0.024). This difference was due to the fact that most responders had an increase of TNFRSF17 of similar magnitude from Day 0 to Day 7 while it was absent in non-responders (Figure 4B).

**Figure 4.**
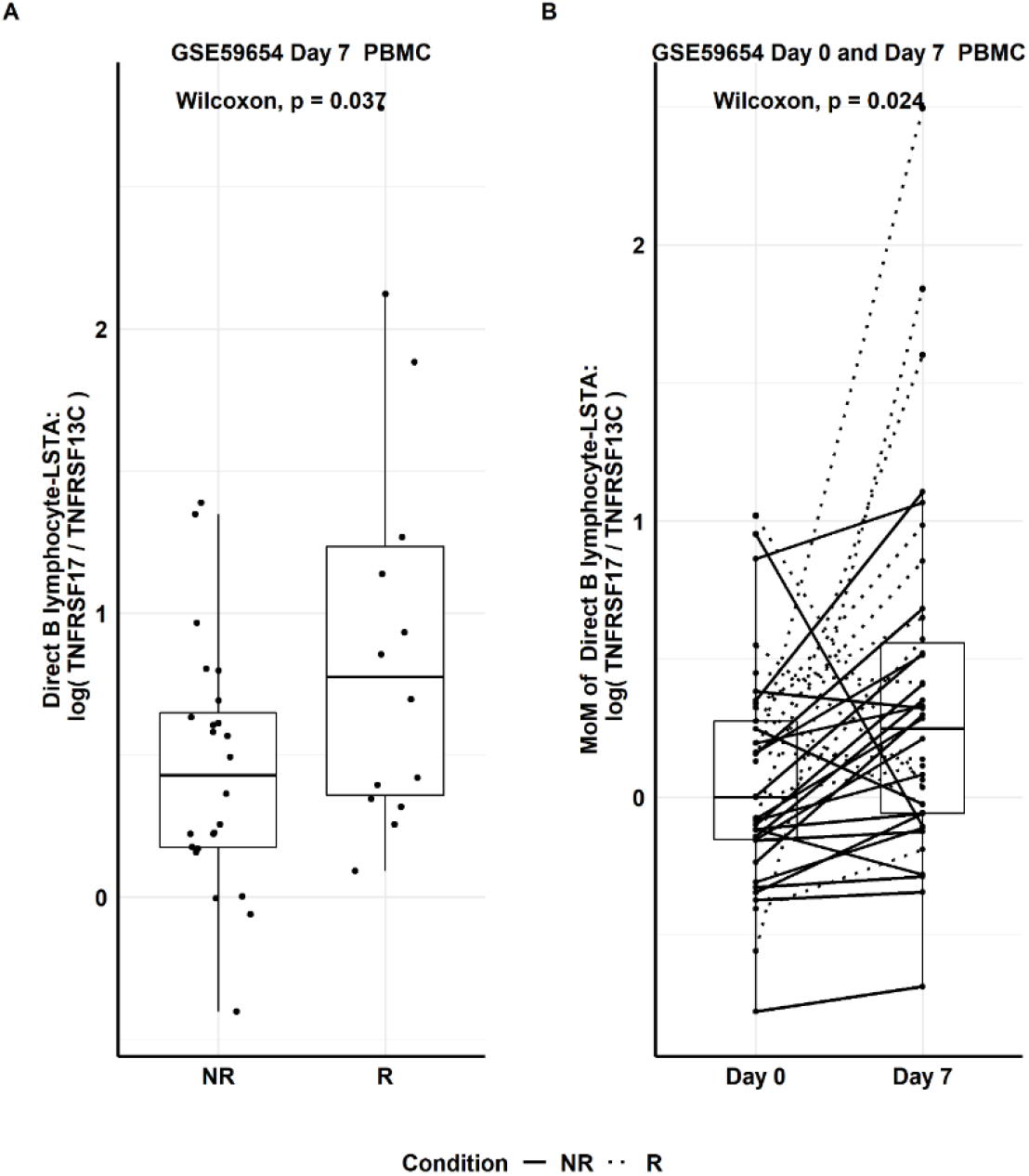
Based on the explorative dataset, GSE59654, (A) Day 7 Direct B-lymphocyte LS-TA results of TNFRSF17 in responder (R) and non-responder (NR) groups. (B) Paired Day 0 and Day 7 change in Direct LS-TA expressed as multiple of median (MoM) using Day 0 group median.

Both the differential Direct B lymphocyte LS-TA of TXNDC5 at Day 7 and the increment from Day 0 to Day 7 were also statistically significant between responders and non-responders (Figure 5).

**Figure 5.**
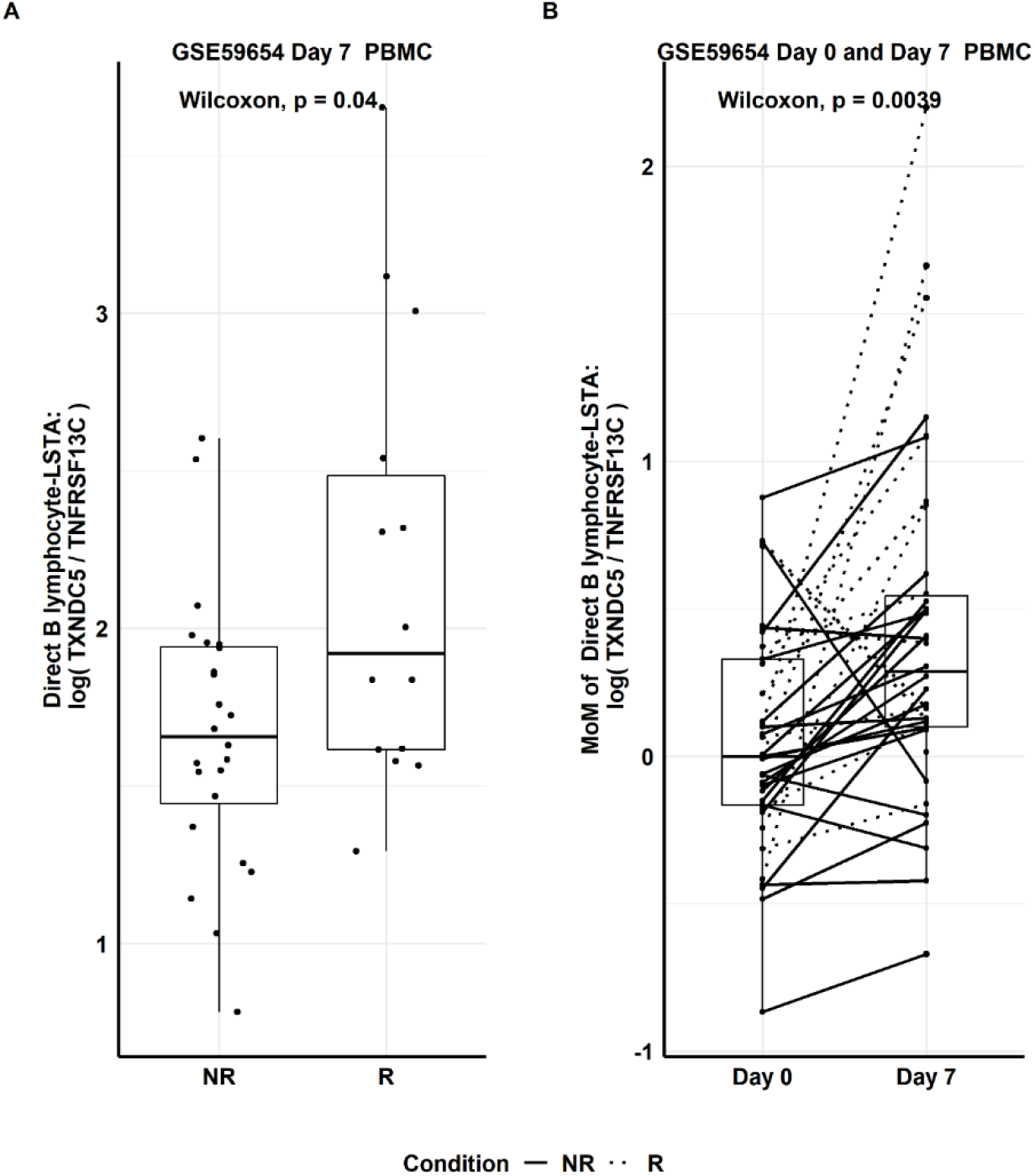
(A) Day 7 Direct B-lymphocyte LS-TA results of another target gene TXNDC5 in responder (R) and non-responder (NR) groups. (B) Paired Day 0 and Day 7 change in Direct LS-TA expressed as multiple of median (MoM) using Day 0 group median.

### Meta-analysis of differentially expressed B lymphocytes LS-TA genes in other datasets

To have a better idea of the reproducibility or replication of these findings, meta-analyses were performed for these 4 Direct B lymphocyte LS-TA biomarkers with PBMC data in a collection of datasets which were performed on different platforms (including Affymetrix and Illumina microarrays).

| Target gene of B lymphocyte | Direct B lymphocyte LS-TA biomarkers using PBMC data:<br>(Ratio of TA of 2 genes as LS-TA biomarker) |
| --- | --- |
| TNFRSF17 | TNFRSF17: TNFRSF13C<br>TNFRSF17: FCRLA |
| TXNDC5 | TXNDC5: TNFRSF13C<br>TXNDC5: FCRLA |

The meta-analysis on TNFRSF17 could be performed on 7 PBMC datasets (Table 1 and Figure 6). Regardless of having TNFRSF13C or FCRLA as the reference gene, the meta-analysis confirmed an overall differential level of Direct LS-TA TNFRSF17 on Day 7 between NR and R groups. In fact, the overall effects were similar in magnitude, and their SMD ranged from 0.6 and 0.8. When TXNDC5 was used as the target gene, similar results were obtained. It suggests that either TNFRSF17 or TXNDC5 and their Direct B lymphocyte LS-TA could be used as a biomarker for vaccination response.

**Figure 6.**
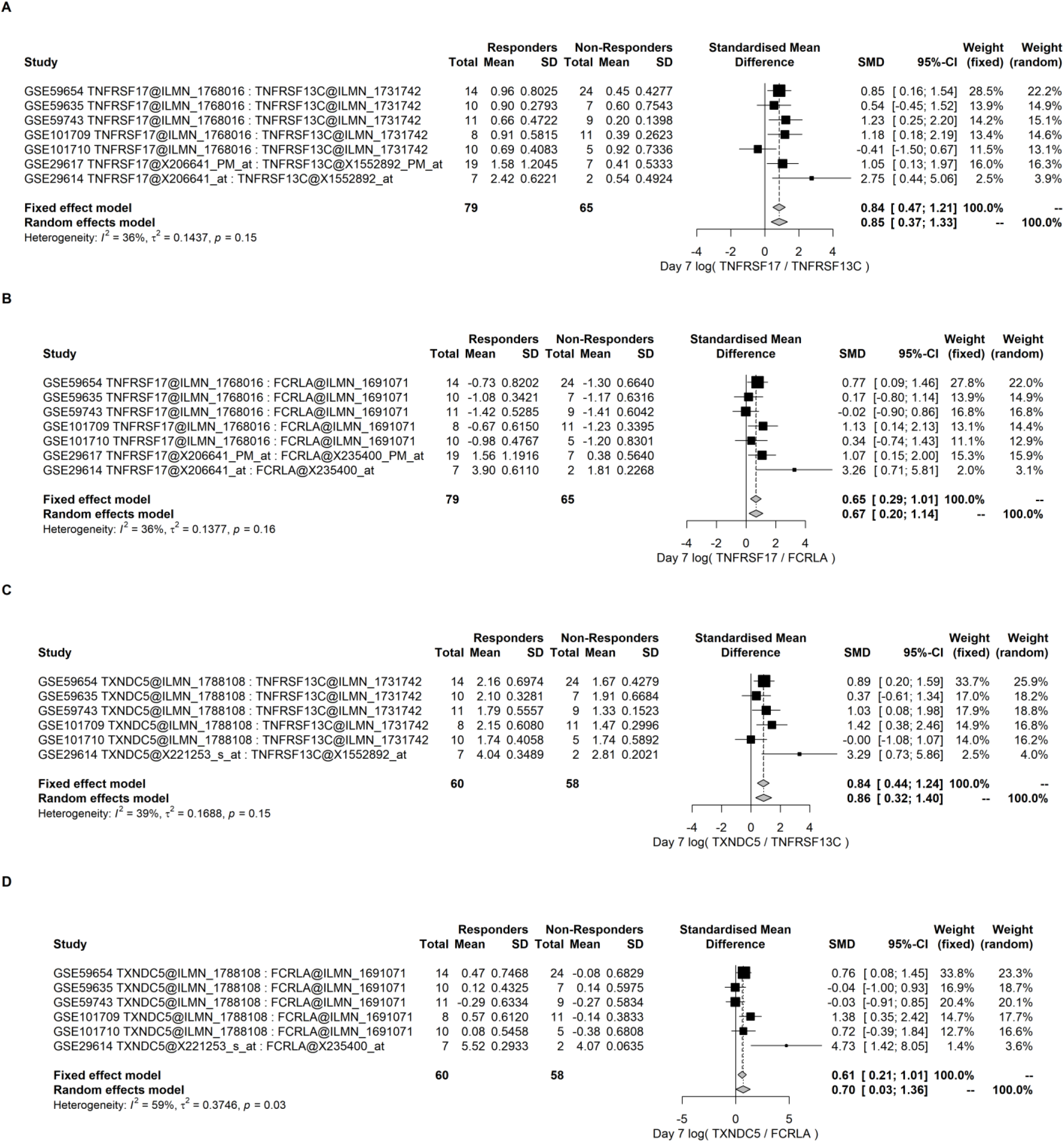
Meta-analysis of the Day7 results of Direct B lymphocyte-LSTA of TNFRSF17 and TXNDC5 using TNFRSF13C or FCRLA as a reference gene in PBMC datasets. Day 7 Direct B lymphocyte-LSTA was significantly higher among the responder groups with an overall effect SMD of 0.6 to 0.8. The specific probe sets used in the calculation of Direct LS-TA are also shown in the table following the gene symbols. (A) and (B) are the results of Direct LS-TA of TNFRSF17 using two different B lymphocyte reference genes (TNFRSF13C and FCRLA). (C) and (D) show the Direct LS-TA results of TXNDC5 using these 2 B lymphocyte reference genes. The specific probe sets used in the calculation of Direct LS-TA are also shown in the table following the gene symbols.

Similarly, the increment from Day 0 to Day 7 was analyzed by meta-analysis in these datasets. The overall effects of greater increment in the responders were confirmed (Figure 7).

**Figure 7.**
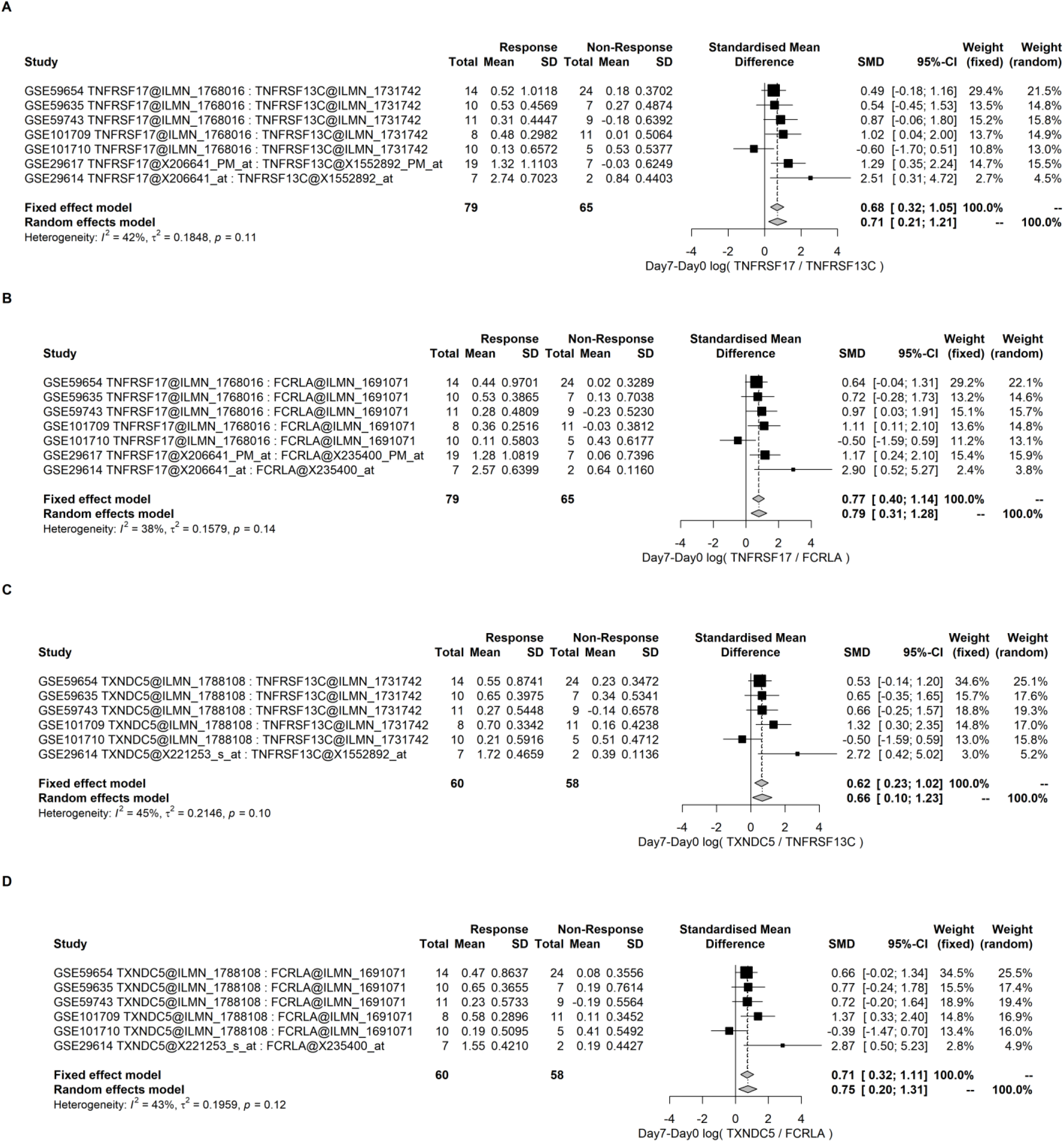
Meta-analysis of the differential increment from Day 0 to Day 7 of Direct B lymphocyte-LSTA between R and NR groups. (A) and (B) are the results of the differential increment of Direct LS-TA of TNFRSF17 using two different B lymphocyte reference genes (TNFRSF13C and FCRLA). (C) and (D) are the results of the differential increment of the Direct LS-TA results of TXNDC5 using these 2 B lymphocyte reference genes. Direct B lymphocyte-LSTA showed significantly higher increments among the responders with an overall effect SMD of between 0.62 to 0.77. The specific probe sets used in the calculation of Direct LS-TA are also shown in the table following the gene symbols.

### Evaluation of the performance by Receiver operator curve (ROC) analysis

Receiver operator curve (ROC) analysis was used to evaluate the test performance of Direct LS-TA as an early (first-week) predictive biomarker for subsequent response status at Day 28 after vaccination. As the study group case-mix were different among studies, the AUC was also different (Figure 7). However, all studies had AUC greater than 0.5, and those of a few studies were higher than 0.8. In some settings, the sensitivity could be 0.8 or higher, while the specificity was around 0.5. On the other hand, a high specificity of the range of 0.8-0.9 could be targeted if a lower sensitivity down to 0.6 is allowed. Figure 8 shows the ROC analysis of the increment of Direct B lymphocyte LS-TA as a biomarker. Again, most studies have AUC above 0.6.

**Figure 8.**
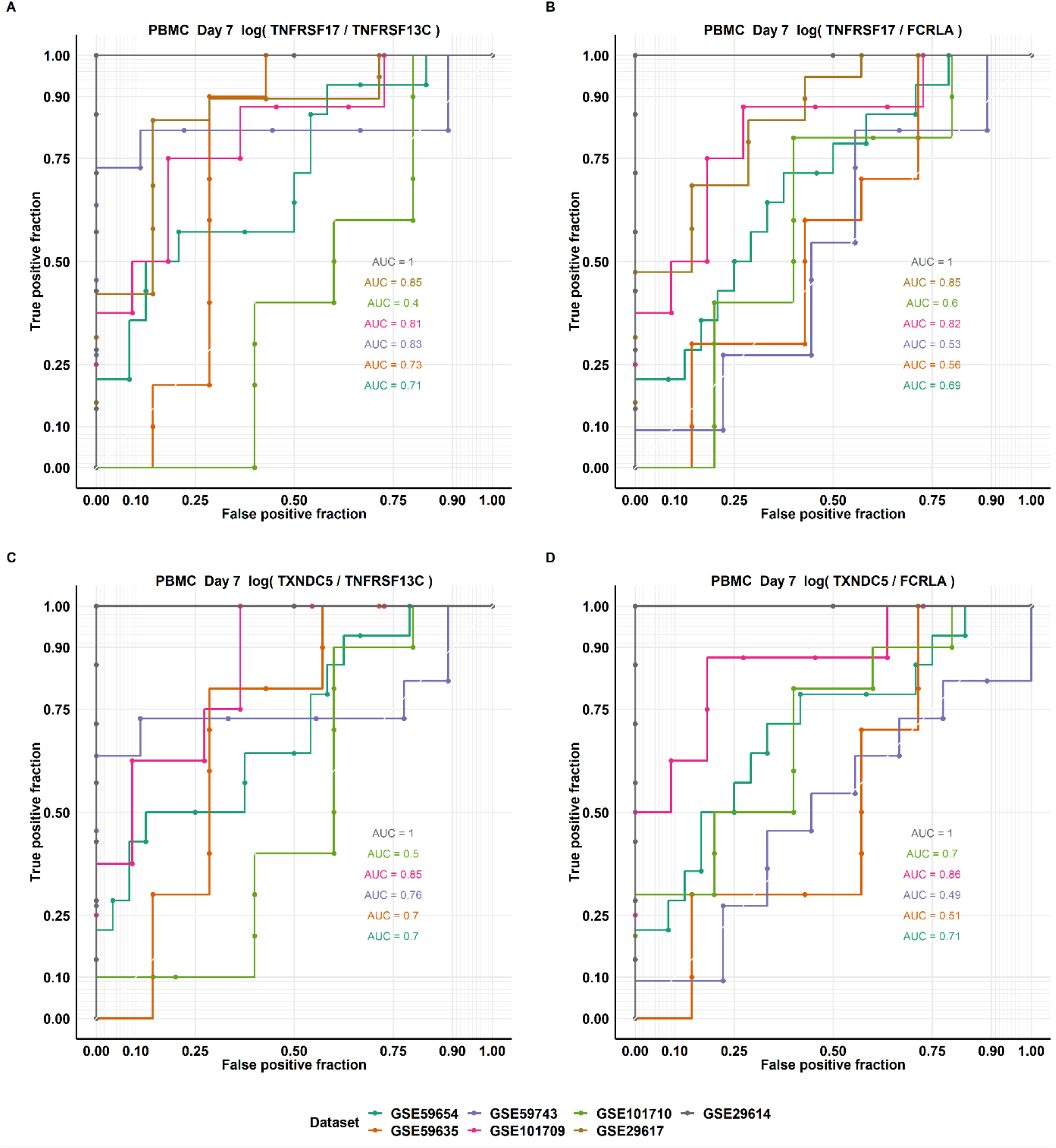
ROC curves of Day 7 Direct B lymphocyte-LSTA of TNFRSF17 and TXNDC5 using two different B lymphocyte informative reference genes (TNFRSF13C and FCRLA). The performance of the single cell-type-specific biomarker parameters Direct LSTA were analyzed by ROC with Responder as a positive outcome.

**Figure 9.**
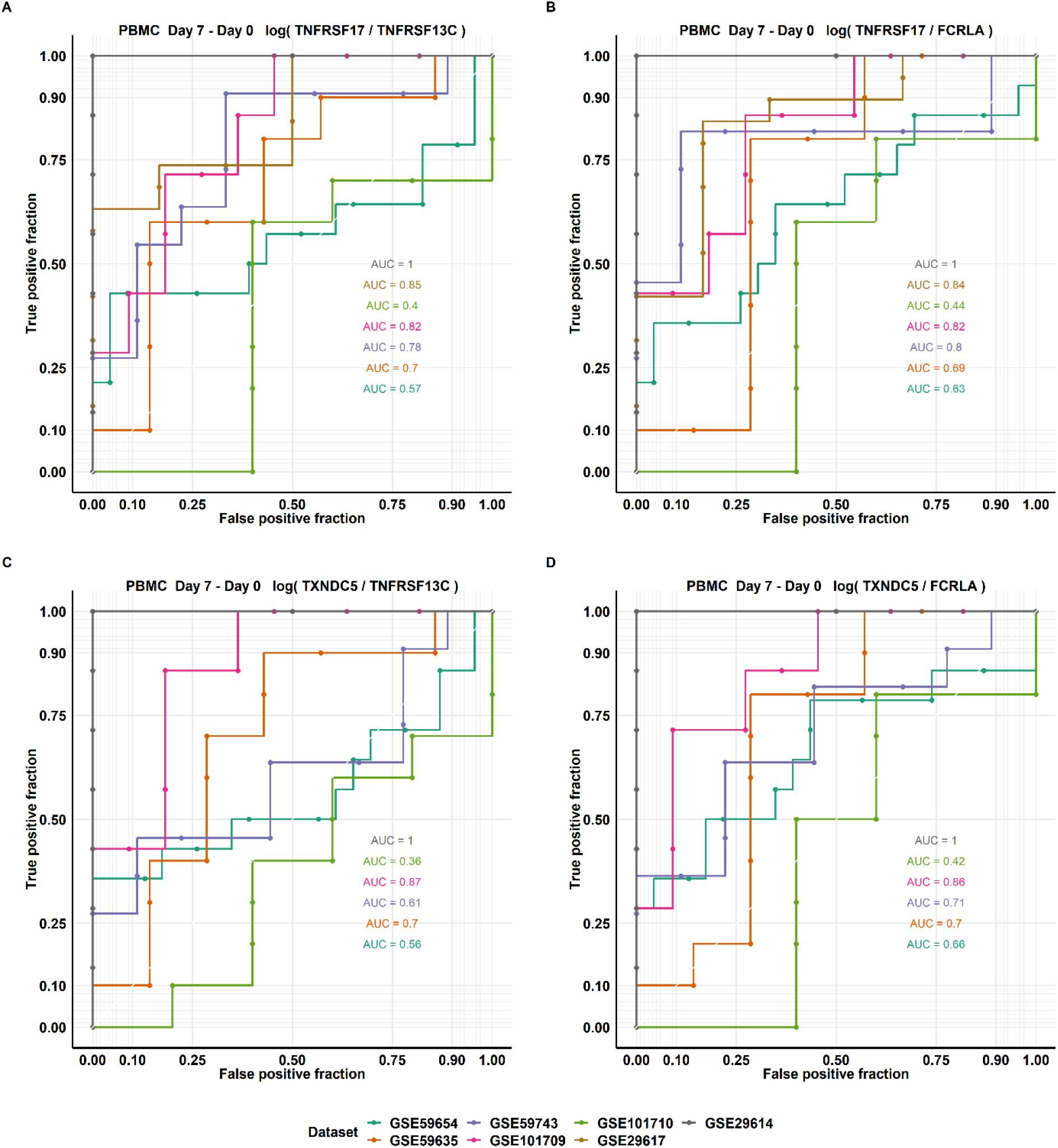
ROC curves of Day 0 to Day 7 increment of Direct B lymphocyte-LSTA of TNFRSF17 and TXNDC5 using two different B lymphocyte informative reference genes (TNFRSF13C and FCRLA). The performance of the single cell-type-specific biomarker parameters Direct LSTA were analyzed by ROC with Responder as a positive outcome.

The exact cut-off point to be used depends on the purpose and rationale of the test. The applications of this method are wide. For example, if the vaccine is unstable, the producer would want to know early if any batch of the vaccine had degraded; this test would provide the answer as early as seven days after administration of the batch. In this scenario, a group of vaccine recipients is monitored for the Day 7 increase in Direct B lymphocyte LS-TA. A high sensitivity cut-off value will be used, so most of the recipients are expected to reach the cut-off value indicating that the vaccine is active. If the number or percentage of recipients showing an increase in Direct B lymphocyte LS-TA on Day 7 is less than a pre-defined percentage which is available on the ROC curve for any specific cut-off value. Degradation of the batch of vaccine would be possible and remedial work could be implemented early rather than waiting for the Day 28 antibody response.

On the other hand, Direct B-lymphocyte LS-TA test could be used for personalized vaccination in which an individual wants to know his subsequent vaccination response early after vaccination.

A high specificity cut-off value will be used in this scenario. Although only around 60% of the responder recipients will achieve this cut-off value, the false position rate is low. This will support a high positive predictive value, so recipients with positive Direct B-lymphocyte LS-TA results would be very likely to be responders. The consequence of a negative Direct B-lymphocyte LS-TA is a requirement of doing a follow-up antibody test on Day 28. The high positive rate of Day 7 Direct B lymphocyte LS-TA will allow the majority of responders to be given the results early.

## Supporting information

Supplementary workflow

## Summary

A simple TA test to determine B lymphocyte gene expression in PBMC samples that could be readily incorporated into the routine clinical laboratory is introduced here. This B lymphocyte Direct LS-TA provides a valid reflection of gene expression of purified B lymphocyte without the need to isolate B lymphocytes from blood cells. It could be used as an early prediction of seroconversion status after influenza vaccination. During the COVID-19 epidemic, a vaccination campaign is just underway. Due to the same mechanism of antibody production, Direct B lymphocyte LS-TA will be a very useful companion assay for the practice of personalized vaccination or as a biological quality control method for monitoring of vaccine efficiency and degradation.

## Conflict of interest statements

Patent application pending.

D Huang and A Liu are employees of Cytomics Ltd.

KS Leung and N Tang are share-holders of Cytomics Ltd.

Cytomics Ltd holds a license to use a patent related to Direct LS-TA assay.

## References

Avey, S., Mohanty, S., Chawla, D. G., Meng, H., Bandaranayake, T., Ueda, I., Zapata, H. J., Park, K., Blevins, T. P., Tsang, S., Belshe, R. B., Kaech, S. M., Shaw, A. C., & Kleinstein, S. H. (2020). Seasonal Variability and Shared Molecular Signatures of Inactivated Influenza Vaccination in Young and Older Adults. Journal of Immunology (Baltimore, Md.: 1950), 204(6), 1661–1673. https://doi.org/10.4049/jimmunol.1900922

Avila Cobos, F., Alquicira-Hernandez, J., Powell, J. E., Mestdagh, P., & De Preter, K. (2020). Benchmarking of cell type deconvolution pipelines for transcriptomics data. Nature Communications, 11(1), 5650. https://doi.org/10.1038/s41467-020-19015-1

Black, S., Bloom, D. E., Kaslow, D. C., Pecetta, S., & Rappuoli, R. (2020). Transforming vaccine development. Seminars in Immunology, 50, 101413. https://doi.org/10.1016/j.smim.2020.101413

Blanco, E., Pérez-Andrés, M., Arriba-Méndez, S., Contreras-Sanfeliciano, T., Criado, I., Pelak, O., Serra-Caetano, A., Romero, A., Puig, N., Remesal, A., Canizales, J. T., López-Granados, E., Kalina, T., Sousa, A. E., Zelm, M. van, Burg, M. van der, Dongen, J. J. M. van, & Orfao, A. (2018). Age-associated distribution of normal B-cell and plasma cell subsets in peripheral blood. Journal of Allergy and Clinical Immunology, 141(6), 2208–2219.e16. https://doi.org/10.1016/j.jaci.2018.02.017

Casey, R. M., Harris, J. B., Ahuka-Mundeke, S., Dixon, M. G., Kizito, G. M., Nsele, P. M., Umutesi, G., Laven, J., Kosoy, O., Paluku, G., Gueye, A. S., Hyde, T. B., Ewetola, R., Sheria, G. K. M., Muyembe-Tamfum, J.-J., & Staples, J. E. (2019). Immunogenicity of Fractional-Dose Vaccine during a Yellow Fever Outbreak—Final Report. The New England Journal of Medicine, 381(5), 444–454. https://doi.org/10.1056/NEJMoa1710430

Committee for Medicinal Products for Human Use. (1997). Note for guidance on harmonisation of requirements for influenza vaccines. European Agency for the Evaluation of Medicinal Products, Brussels, Belgium (1997). https://www.ema.europa.eu/en/documents/scientific-guideline/note-guidance-harmonisation-requirements-influenza-vaccines_en.pdf

Ding, Y., Zhou, L., Xia, Y., Wang, W., Wang, Y., Li, L., Qi, Z., Zhong, L., Sun, J., Tang, W., Liang, F., Xiao, H., Qin, T., Luo, Y., Zhao, X., Shu, Z., Ru, Y., Dai, R., Wang, H., … Zhao, X. (2018). Reference values for peripheral blood lymphocyte subsets of healthy children in China. Journal of Allergy and Clinical Immunology, 142(3), 970–973.e8. https://doi.org/10.1016/j.jaci.2018.04.022

Henn, A. D., Wu, S., Qiu, X., Ruda, M., Stover, M., Yang, H., Liu, Z., Welle, S. L., Holden-Wiltse, J., Wu, H., & Zand, M. S. (2013). High-Resolution Temporal Response Patterns to Influenza Vaccine Reveal a Distinct Human Plasma Cell Gene Signature. Scientific Reports, 3(1), 2327. https://doi.org/10.1038/srep02327

Horns, F., Dekker, C. L., & Quake, S. R. (2020). Memory B Cell Activation, Broad Anti-influenza Antibodies, and Bystander Activation Revealed by Single-Cell Transcriptomics. Cell Reports, 30(3), 905–913.e6. https://doi.org/10.1016/j.celrep.2019.12.063

Mahalanobis distance. (2020). In Wikipedia. https://en.wikipedia.org/w/index.php?title=Mahalanobis_distance&oldid=995007639

Mo, Z., Nong, Y., Liu, S., Shao, M., Liao, X., Go, K., & Lavis, N. (2017). Immunogenicity and safety of a trivalent inactivated influenza vaccine produced in Shenzhen, China. Human Vaccines & Immunotherapeutics, 13(6), 1272–1278. https://doi.org/10.1080/21645515.2017.1285475

Monaco, G., Lee, B., Xu, W., Mustafah, S., Hwang, Y. Y., Carré, C., Burdin, N., Visan, L., Ceccarelli, M., Poidinger, M., Zippelius, A., Pedro de Magalhães, J., & Larbi, A. (2019). RNA-Seq Signatures Normalized by mRNA Abundance Allow Absolute Deconvolution of Human Immune Cell Types. Cell Reports, 26(6), 1627–1640.e7. https://doi.org/10.1016/j.celrep.2019.01.041

Nakaya, H. I., Wrammert, J., Lee, E. K., Racioppi, L., Marie-Kunze, S., Haining, W. N., Means, A. R., Kasturi, S. P., Khan, N., Li, G.-M., McCausland, M., Kanchan, V., Kokko, K. E., Li, S., Elbein, R., Mehta, A. K., Aderem, A., Subbarao, K., Ahmed, R., & Pulendran, B. (2011). Systems biology of vaccination for seasonal influenza in humans. Nature Immunology, 12(8), 786–795. https://doi.org/10.1038/ni.2067

Pezeshki, A., Ovsyannikova, I. G., McKinney, B. A., Poland, G. A., & Kennedy, R. B. (2019). The role of systems biology approaches in determining molecular signatures for the development of more effective vaccines. Expert Review of Vaccines, 18(3), 253–267. https://doi.org/10.1080/14760584.2019.1575208

Rogers, L. R. K., de los Campos, G., & Mias, G. I. (2019). Microarray Gene Expression Dataset Re-analysis Reveals Variability in Influenza Infection and Vaccination. Frontiers in Immunology, 10. https://doi.org/10.3389/fimmu.2019.02616

Shen-Orr, S. S., & Gaujoux, R. (2013). Computational deconvolution: Extracting cell type-specific information from heterogeneous samples. Current Opinion in Immunology, 25(5), 571–578. https://doi.org/10.1016/j.coi.2013.09.015

Shen-Orr, S. S., Tibshirani, R., Khatri, P., Bodian, D. L., Staedtler, F., Perry, N. M., Hastie, T., Sarwal, M. M., Davis, M. M., & Butte, A. J. (2010). Cell type-specific gene expression differences in complex tissues. Nature Methods, 7(4), 287–289. https://doi.org/10.1038/nmeth.1439

Thakar, J., Mohanty, S., West, A. P., Joshi, S. R., Ueda, I., Wilson, J., Meng, H., Blevins, T. P., Tsang, S., Trentalange, M., Siconolfi, B., Park, K., Gill, T. M., Belshe, R. B., Kaech, S. M., Shadel, G. S., Kleinstein, S. H., & Shaw, A. C. (2015). Aging-dependent alterations in gene expression and a mitochondrial signature of responsiveness to human influenza vaccination. Aging, 7(1), 38–52. https://doi.org/10.18632/aging.100720

Tsang, J. S., Schwartzberg, P. L., Kotliarov, Y., Biancotto, A., Xie, Z., Germain, R. N., Wang, E., Olnes, M. J., Narayanan, M., Golding, H., Moir, S., Dickler, H. B., Perl, S., Cheung, F., Baylor HIPC Center, & CHI Consortium. (2014). Global analyses of human immune variation reveal baseline predictors of postvaccination responses. Cell, 157(2), 499–513. https://doi.org/10.1016/j.cell.2014.03.031

Uhlen, M., Karlsson, M. J., Zhong, W., Tebani, A., Pou, C., Mikes, J., Lakshmikanth, T., Forsström, B., Edfors, F., Odeberg, J., Mardinoglu, A., Zhang, C., von Feilitzen, K., Mulder, J., Sjöstedt, E., Hober, A., Oksvold, P., Zwahlen, M., Ponten, F., … Brodin, P. (2019). A genome-wide transcriptomic analysis of protein-coding genes in human blood cells. Science (New York, N.Y.), 366(6472). https://doi.org/10.1126/science.aax9198

Weiner, J., Lewis, D. J. M., Maertzdorf, J., Mollenkopf, H.-J., Bodinham, C., Pizzoferro, K., Linley, C., Greenwood, A., Mantovani, A., Bottazzi, B., Denoel, P., Leroux-Roels, G., Kester, K. E., Jonsdottir, I., van den Berg, R., Kaufmann, S. H. E., & Del Giudice, G. (2019). Characterization of potential biomarkers of reactogenicity of licensed antiviral vaccines: Randomized controlled clinical trials conducted by the BIOVACSAFE consortium. Scientific Reports, 9(1), 20362. https://doi.org/10.1038/s41598-019-56994-8

