## Supplementary figures and images for "Direct measurement of B lymphocyte gene expression biomarkers in peripheral blood enables early prediction of seroconversion after vaccination"

### Supplementary Figure1

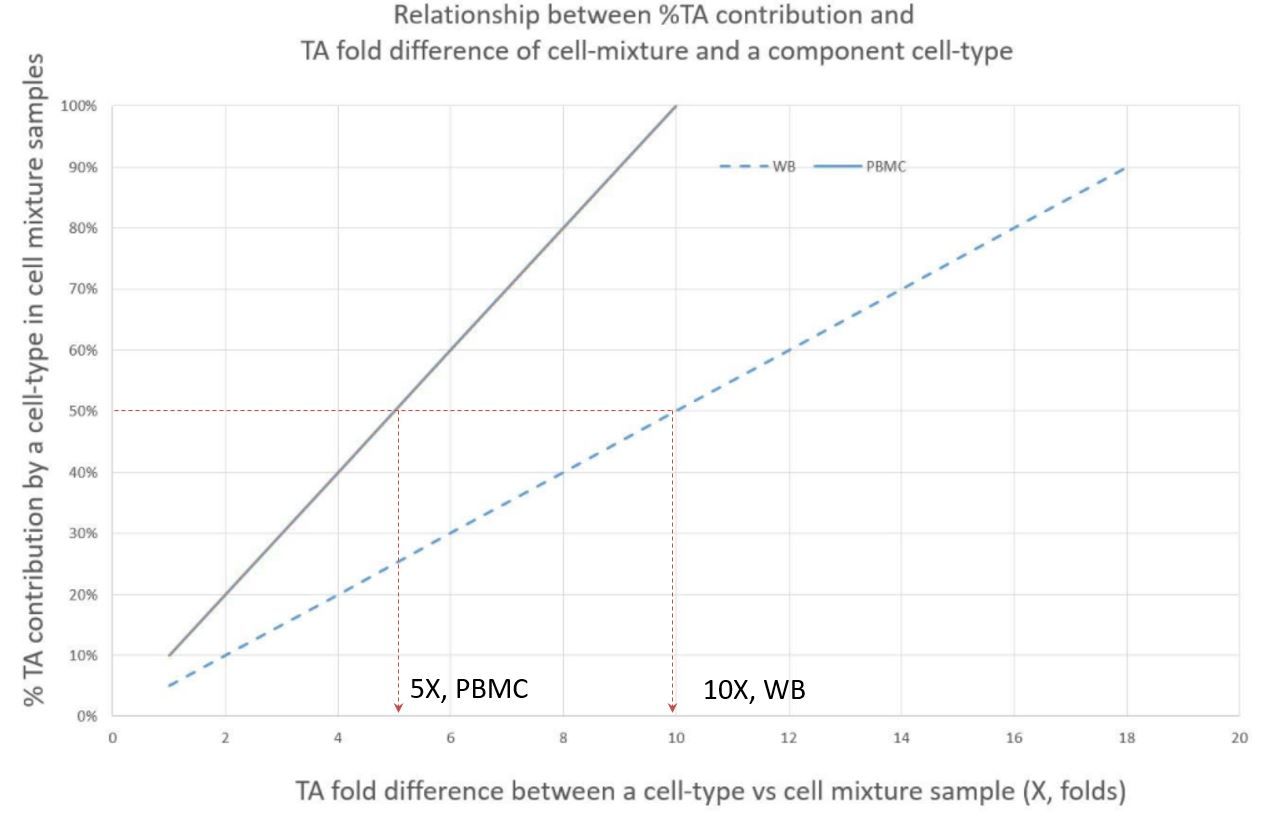

### Supplementary workflow

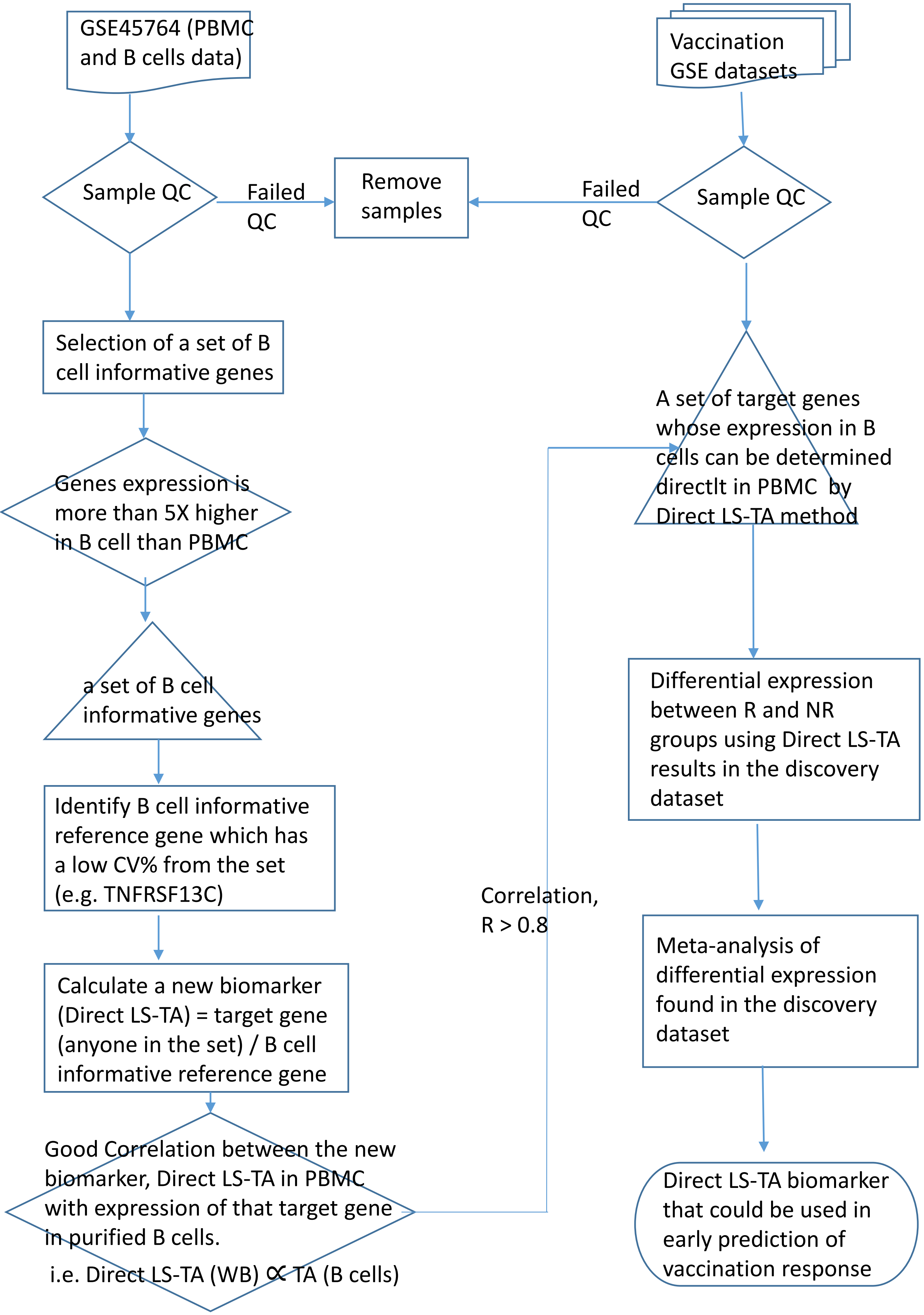
