## Supplementary table1 for "Direct measurement of B lymphocyte gene expression biomarkers in peripheral blood enables early prediction of seroconversion after vaccination"

| Supplementary table1 Shortlisted potential B lymphocyte informative genes in RNA-seq dataset GSE45764 | | | |
| --- | --- | --- | --- |
| Gene symbol | Expression fold change between purified B lymphocytes and cell mixture samples | Median Log2 expression in purified B lymphocyte samples | CV% in purified B lymphocyte samples |
| RASGRP3 | 11.790 | 7.310 | 0.09 |
| COBLL1 | 12.820 | 7.550 | 0.1 |
| HLA-DOB | 13.450 | 8.300 | 0.1 |
| BLNK | 10.780 | 7.370 | 0.1 |
| RALGPS2 | 12.470 | 9.870 | 0.15 |
| EBF1 | 16.110 | 6.230 | 0.15 |
| CD19 | 15.240 | 8.490 | 0.15 |
| CD79B | 11 | 9.590 | 0.15 |
| CD79A | 15.780 | 10.400 | 0.15 |
| LARGE1 | 13.640 | 5.720 | 0.15 |
| OSBPL10 | 16.340 | 7.860 | 0.16 |
| BANK1 | 15.350 | 9.900 | 0.16 |
| PNOC | 11.550 | 5.710 | 0.16 |
| PAX5 | 15.780 | 8.730 | 0.16 |
| POU2AF1 | 14.220 | 8.990 | 0.16 |
| TNFRSF13C | 14.620 | 6.980 | 0.16 |
| TLR10 | 12.040 | 8.410 | 0.17 |
| SYNPO | 16.910 | 6.910 | 0.17 |
| MS4A1 | 16.220 | 11.810 | 0.17 |
| FAM30A | 14.220 | 9.060 | 0.17 |
| P2RX5 | 11 | 7.960 | 0.17 |
| BLK | 16.110 | 9.160 | 0.18 |
| ADAM28 | 10.270 | 8.140 | 0.18 |
| CD22 | 16 | 10.430 | 0.18 |
| ZNF860 | 15.560 | 5.920 | 0.19 |
| BEND4 | 10.630 | 5.610 | 0.19 |
| STAP1 | 10.130 | 7.110 | 0.19 |
| DENND5B | 13.550 | 7.520 | 0.19 |
| PCDH9 | 16.220 | 6.240 | 0.19 |
| FCRL2 | 14.930 | 8.550 | 0.2 |
| FCRLA | 14.930 | 8.580 | 0.2 |
| PLEKHG1 | 12.550 | 7.320 | 0.2 |
| CLEC17A | 12.730 | 6.910 | 0.2 |
| VPREB3 | 16 | 6.110 | 0.2 |
| SNX22 | 14.320 | 7.570 | 0.21 |
| LOC284749 | 15.240 | 6.080 | 0.21 |
| FCRL1 | 15.240 | 9.990 | 0.22 |
| CPNE5 | 13.740 | 8.330 | 0.22 |
| ABCB4 | 13.090 | 5.850 | 0.22 |
| CD72 | 10.630 | 7.520 | 0.22 |
| LAMA5 | 13.270 | 7.010 | 0.22 |
| BACE2 | 11.960 | 5.940 | 0.22 |
| SCN3A | 15.780 | 6.160 | 0.23 |
| LCN10 | 14.120 | 6.290 | 0.23 |
| KLHL14 | 14.720 | 6.730 | 0.23 |
| NIBAN3 | 11.960 | 9.370 | 0.23 |
| KCNH8 | 14.420 | 5.800 | 0.25 |
| BHLHE41 | 14.930 | 5.480 | 0.25 |
| FCER2 | 14.320 | 8.330 | 0.25 |
| COL4A4 | 12.910 | 6.380 | 0.26 |
| COL4A3 | 16.910 | 6.980 | 0.27 |
| LOC283663 | 13.830 | 9.760 | 0.27 |
| TNFRSF13B | 17.030 | 6.500 | 0.27 |
| PKIG | 11.160 | 6.210 | 0.27 |
| LAMC1 | 10.130 | 6.620 | 0.28 |
| CNTNAP2 | 17.270 | 6.370 | 0.29 |
| MACROD2 | 14.930 | 6.080 | 0.29 |
| PARM1 | 10.780 | 6.380 | 0.3 |
| COL19A1 | 14.220 | 6.940 | 0.31 |
| SSPN | 14.030 | 5.460 | 0.31 |
| TCL1A | 13.360 | 8.740 | 0.31 |
| CLCN4 | 10.270 | 5.800 | 0.32 |
| CD200 | 16.560 | 6.730 | 0.33 |
| FCRL5 | 16.340 | 9.020 | 0.35 |
| IGKC | 15.670 | 12.550 | 0.36 |
| EML6 | 14.520 | 6.900 | 0.38 |
| PEG10 | 12.040 | 5.460 | 0.38 |
| IGHD | 11.310 | 8.450 | 0.4 |
| COCH | 15.030 | 6.360 | 0.44 |
| TNFRSF17 | 12.470 | 6.280 | 0.62 |
| JUN | 13.450 | 8.240 | 0.63 |
| IGHG2 | 15.450 | 8.570 | 0.63 |
| JCHAIN | 11.390 | 11.610 | 0.7 |
| MZB1 | 10.850 | 7.790 | 0.79 |
| IGHG3 | 13.930 | 7.630 | 0.81 |
| IGHM | 15.780 | 11.010 | 0.91 |
| IGHG4 | 17.750 | 5.520 | 0.96 |
| IGHG1 | 12.550 | 11.350 | 0.99 |
| IGHA2 | 13.550 | 7.820 | 1.02 |
| TXNDC5 | 10.560 | 5.320 | 1.03 |
| DSP | 16.450 | 5.840 | 1.05 |
| IGHA1 | 14.720 | 11.410 | 1.05 |
| IGLL5 | 19.430 | 10.390 | 1.14 |
