## Supplementary material for "Direct measurement of B lymphocyte gene expression biomarkers in peripheral blood enables early prediction of seroconversion after vaccination": Table1

Table 1. List of gene expression datasets of PBMC or WB used in this study

| Data series accession number | Type of Blood samples(WB or PBMC) | Type of vaccine used (against the virus) | Total No. of samples (all time points) | No.of QC failed samples | No.of Non-Responders (NR) at Day7 | No.of Responders (R ) at Day7 | Reference |
| --- | --- | --- | --- | --- | --- | --- | --- |
| GSE29614 | PBMC | Influenza TIV | 18 | - | 2 | 7 | (Nakaya et al., 2011) |
| GSE29615 | PBMC | Influenza LAIV | 55 | - | 26 | 1 | (Nakaya et al., 2011) |
| GSE29617 | PBMC | Influenza TIV | 53 | 2 | 7 | 19 | (Nakaya et al., 2011) |
| GSE101709 | PBMC | Influenza | 41 | 4 | 11 | 8 | (Avey et al., 2020) |
| GSE101710 | PBMC | Influenza | 31 | - | 5 | 10 | (Avey et al., 2020) |
| GSE59635 | PBMC | Influenza | 36 | 1 | 7 | 10 | (Thakar et al., 2015) |
| GSE59654 | PBMC | Influenza | 76 | 3 | 24 | 14 | (Thakar et al., 2015) |
| GSE59743 | PBMC | Influenza | 50 | - | 9 | 11 | (Thakar et al., 2015) |
| GSE4576  4 | Paired PBMC and Purified B lymphocytes | Influenza | 104 | 6 |  |  | (Henn et al., 2013) |
