## Supplementary material for "Direct measurement of B lymphocyte gene expression biomarkers in peripheral blood enables early prediction of seroconversion after vaccination": Table2

Table 2. Shortlisted potential B lymphocyte informative genes in RNA-seq dataset GSE45764. Cell-type informative reference genes could be identified among those with low CV%. Two genes stand out among peers of other well-recognized B cell markers, TNFRSF13C and FCRLA, which are further analyzed as B cell informative reference genes in this manuscript.

| Gene symbol | Expression fold change between purified B lymphocytes and WB samples (X, folds) | CV% in purified B lymphocyte samples |
| --- | --- | --- |
| *HLA-DOB* | 13.360 | 11% |
| *CD19* | 15.140 | 15% |
| *CD79B* | 10.930 | 15% |
| *CD79A* | 15.560 | 15% |
| *BANK1* | 15.670 | 16% |
| ***TNFRSF13C*** | 14.420 | 16% |
| *MS4A1* | 16 | 17% |
| *BLK* | 16 | 18% |
| *CD22* | 16.340 | 18% |
| ***FCRLA*** | 14.830 | 20% |
| *FCER2* | 14.220 | 24% |
| *TNFRSF13B* | 16.910 | 26% |
| *TCL1A* | 13.550 | 30% |
| *TNFRSF17* | 12.640 | 62% |
| *JCHAIN* | 11.390 | 70% |
| *TXNDC5* | 10.560 | 103% |
