## Supplementary table2 for "Direct measurement of B lymphocyte gene expression biomarkers in peripheral blood enables early prediction of seroconversion after vaccination"

|  |  |  |
| --- | --- | --- |
| Supplement table2 B-lymphocyte target genes that can be reliably measured in PBMC without cell purification | | |
| B lymphocyte informative target gene | B lymphocyte informative reference gene | Pearson��s correlation coefficient between PBMC and B cell |
| IGHG1 | FCRLA | 0.97 |
| IGHG3 | FCRLA | 0.95 |
| MZB1 | FCRLA | 0.94 |
| DSP | FCRLA | 0.94 |
| IGHA1 | FCRLA | 0.94 |
| IGLL5 | FCRLA | 0.94 |
| FCRL5 | FCRLA | 0.93 |
| JCHAIN | FCRLA | 0.93 |
| TNFRSF17 | FCRLA | 0.93 |
| TXNDC5 | FCRLA | 0.91 |
| IGHA2 | FCRLA | 0.91 |
| IGHM | FCRLA | 0.91 |
| IGKC | FCRLA | 0.9 |
| PARM1 | FCRLA | 0.88 |
| COCH | FCRLA | 0.88 |
| IGHG2 | FCRLA | 0.87 |
